# Tissue retention of dendritic cells drives age-associated protective tolerogenic lymphoid structures

**DOI:** 10.64898/2026.10.04.756474

**Authors:** Ines Marin, Blanca M. Morales, Brian Orcutt-Jahns, Jessica Preston, Kevin Marroquin, Zhichang Yang, Ying Zhu, Joshua Webster, Christine Moussion

## Abstract

Ectopic lymphoid structures are typically associated with chronic tissue inflammation in autoimmunity and cancer, where they can amplify local immune responses. Similar structures arise spontaneously with age, yet their origin and function remain unclear. Here we identify tissue retention of mature migratory conventional dendritic cells (cDCs) as a driver of age-associated lymphoid organization. Mature cDCs accumulated within lymphoid structures in aged human and mouse tissues. We demonstrate that impaired cDC egress or pharmacological cDC expansion induced de novo structures in mice across multiple otherwise homeostatic organs without associated pathology. Strikingly, senescence-associated tissue cues, which promoted a distinct cDC maturation state, were required for lymphoid niche initiation. Unlike inflammatory structures described in disease, these cDC-driven structures preferentially accumulated tissue-adapted regulatory T cells and protected against inflammatory injury, revealing a previously unrecognized regulatory dimension of age-associated lymphoid organization.

## Main

In response to prolonged stress or inflammation, tissues recruit immune cells to form localized clusters, known as ectopic lymphoid aggregates that can progressively organize into tertiary lymphoid structures (TLS)^1–3^. Acting as functional niches that mirror secondary lymphoid organs, lymphoid structures can orchestrate adaptive immunity locally and shape disease trajectories across cancer, infection, and autoimmunity^1,4^. The functional consequences of their presence in tissues, however, vary markedly with context: TLS can support protective immunity in cancer and infection^4–6^, sustain pathogenic responses in autoimmune and inflammatory diseases^1,7^, and contribute to mucosal homeostasis at barrier tissues^8,9^. Lymphoid aggregates and TLS-like structures also emerge spontaneously in tissues with aging^10–12^, although evidence remains anecdotic and largely tissue-specific, and their biology during physiological aging is poorly defined. The cellular mechanisms that initiate and sustain lymphoid organization across aged tissues as well as their functional character are still unclear.

Aging and chronic tissue stress profoundly remodel both immune and tissue biology. Hallmarks of this remodeling include myeloid-skewed hematopoiesis, deterioration of tissue lymphatic drainage and the accumulation of senescent cells, which promote tissue ‘inflammaging’ through the continuous release of senescence-associated secretory factors^13–18^. Conventional dendritic cells (cDC) are crucial myeloid cells that sense tissue-derived signals, acquire a migratory program characterized by CCR7 expression, and traffic from the tissue to the draining lymph node (LN), where they orchestrate adaptive immunity and tolerance^19,20^. With age, intrinsic CCR7-dependent cDC migratory capacity declines by approximately half^21,22^, which together with lymphatic dysfunction, restrict tissue cDC migration to the draining LN. How tissue cDCs contribute to tissue immune organization during aging and shape its functional character is largely underexplored.

### Mature cDCs mark lymphoid niches

To characterize the conventional dendritic cell (cDC) compartment in aged tissues, we first analyzed a published single-cell RNA-sequencing dataset from human lung across ages^23^. The most prominent compositional change was the emergence of a mature cDC population in aged tissues minimally represented in young tissues (Fig. 1a, Extended Data Fig. 1a). These cells expressed a conserved cDC maturation program^19^ marked by the expression of *LAMP3*, coestimulatory genes *CD80, CD86,* and *CD83,* and a classical migratory program characterized by *CCR7* and *FSCN1* expression (Extended Data Fig. 1b).

**Fig. 1.**
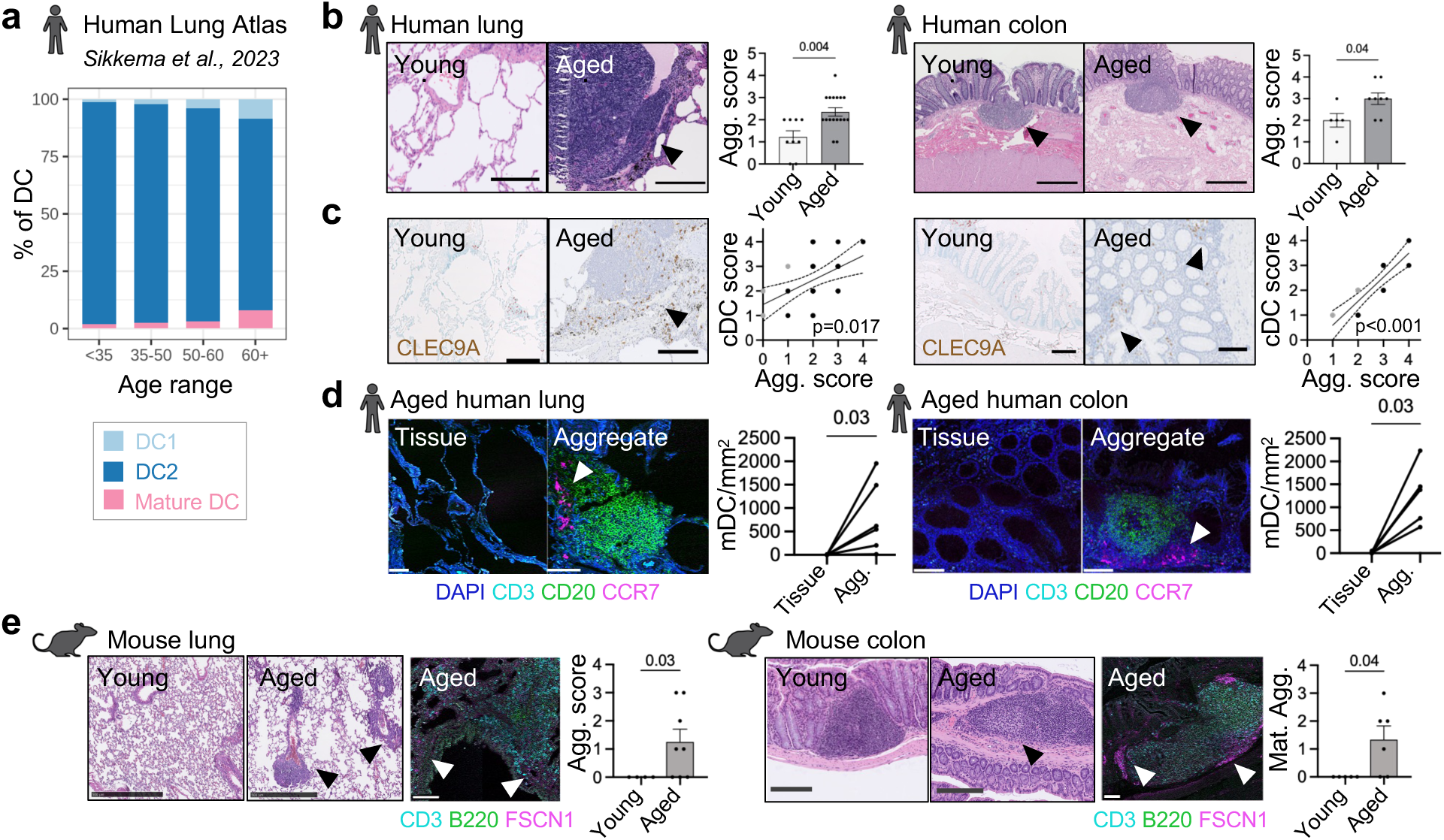
Lymphoid aggregates in aged tissues are enriched in mature cDCs. a, Proportion of conventional dendritic cells (cDCs) in human lung from public single-cell RNA-sequencing dataset^23^. **b,** Representative hematoxylin & eosin (H&E) images and lymphoid-aggregate scores (Agg. score) in lung and colon from healthy individuals: young (<50 years, *n* = 9 lung, 5 colon) and aged (>70 years, *n* = 17 lung, 8 colon) presenting lymphoid aggregates. Two-tailed unpaired *t*-test. Scale bars, 300 µm (lung) and 500 µm (colon). **c,** Representative CLEC9A immunohistochemistry and correlation between CLEC9A+ cDC and lymphoid-aggregate scores in young (gray points; *n* = 9 lung, 5 colon) and aged (black points; *n* = 17 lung, 8 colon) tissues presenting lymphoid aggregates. Linear regression and two-tailed Spearman correlation (r = 0.4646 lung, 0.8631 colon). Scale bars, 300 µm (lung) and 200 µm (colon). **d,** Representative immunofluorescence of CD3+ T cells, CD20+ B cells and CCR7+CD20−CD3− mature cDCs (mDCs) in parenchyma and lymphoid aggregates from aged tissues, with paired mDC quantification between compartments (*n* = 6 lung, 7 colon). Two-tailed paired *t*-test. Scale bars, 100 µm (lung) and 500 µm (colon). **e,** Representative H&E images of lung and colon from young (<4 months; *n* = 4 lung, 5 colon) and aged (>18 months; *n* = 8 lung, 6 colon) mice and immunofluorescence of CD3+ T cells, B220+ B cells and FSCN1+ mature cDCs in lymphoid aggregates. Aggregate score in lung and burden of mature aggregates (Mat. Agg.) containing germinal centres per colon section were quantified. Two-tailed unpaired *t*-test. Scale bars, 500 µm (lung), 200 µm (colon) and 100 µm (immunofluorescence). Data are mean ± s.e.m.

We then examined young and aged human tissues spanning a range of lymphoid aggregate burden (Fig. 1b and Extended Data Fig. 1c). In lung, colon, and liver tissues, lymphoid aggregates were enriched in aged tissues in comparison to young tissues (Fig. 1b and Extended Data Fig. 1c). While in young tissues CLEC9A^+^ cDCs were rather sparse and distributed throughout the tissue parenchyma, in aged tissues, cDCs were more abundant and preferentially clustered within lymphoid aggregates (Fig. 1c and Extended Data Fig. 1d). Consistent with this, increased lymphoid aggregate presence was accompanied by greater cDC abundance in tissues (Fig. 1b,c and Extended Data Fig. 1d). In line with the mature cDC state identified transcriptionally, CCR7^+^ and LAMP3^+^ mature cDCs were mostly exclusively present within lymphoid aggregates and were virtually absent in the surrounding tissue parenchyma (Fig. 1d and Extended Data Fig. 1f). These lymphoid aggregates represented organized structures with B cell and T cell zones, and mature cDCs preferentially accumulated in T cell-rich areas (Fig. 1d and Extended Data Fig. 1f). Together, these spatial analyses show that, rather than migrating to draining lymph nodes (LNs), certain CCR7^+^ mature migratory cDCs might preferentially accumulate within ectopic lymphoid aggregates within aged tissues.

Because cross-sectional human data cannot resolve whether these structures arise as a feature of aging itself or in response to chronic infection or environmental exposures, we next examined their occurrence in a genetically and environmentally controlled preclinical aging mouse model. Aged mice developed spontaneously similar ectopic lymphoid aggregates in multiple otherwise homeostatic tissues including lung, salivary gland, liver, and pancreas, and showed increased maturation of colonic lymphoid structures, harboring evident germinal centers (Fig. 1e and Extended Data Fig. 1g). Similarly, these aged mice lymphoid structures were enriched in FSCN1⁺ mature migratory cDCs (Fig. 1e).

Together, these observations identify the spontaneous development of organized lymphoid structures with an accumulation of mature migratory cDCs as a recurrent structural feature of lymphoid niches across aging tissues in mouse and human.

### cDC tissue retention initiates lymphoid niches

The observed association between mature cDC accumulation and lymphoid structure development in aged tissues cannot distinguish whether mature cDC accumulation contributes to lymphoid organization or instead results from it. We therefore asked whether blocking the egress of mature cDCs from otherwise homeostatic tissues is sufficient to initiate ectopic lymphoid niche formation. Upon maturation, cDCs normally upregulate the chemokine receptor CCR7 and migrate to tissue-draining LNs following the gradients of CCR7 ligands^20^. To induce the retention of mature cDC in tissues, we generated mice lacking CCR7 in the cDC lineage^24^ (*Clec9a^cre^Ccr7^fl/fl^,* hereafter CCR7^ΔcDC^; Figure 2a). CCR7 deletion in CCR7^ΔcDC^ mice was restricted to cDCs and spared T and B lymphocytes, as confirmed by locus recombination analysis in sorted immune populations and lineage tracing mouse models of Cre expression (*Clec9a^cre^CCR7^fl/fl^Rosa26^LSL.tdtomato^*) (Extended Data Fig. 2a,b). CCR7^ΔcDC^ cDCs (both cDCs type 1 and 2) lacked CCR7 expression upon LPS-induced maturation (Extended Data 2c) and presented a lack of CCR7-dependent migratory capacity measured in an in vivo DC migration assay, whereas other DC populations, including Langerhans cells, monocyte-derived DC and plasmacytoid DCs, retained migration to the LNs (Extended Data Fig. 2d,e). Accordingly, CCR7^ΔcDC^ mice showed reduced proportion of migratory cDCs in tissue-draining LNs (Figure 2b and Extended Data Fig. 2f) and increased burden of cDCs in peripheral tissues when compared to CCR7^wt^ control mice (Fig. 2c and Extended Data Fig. 2g).

**Fig. 2.**
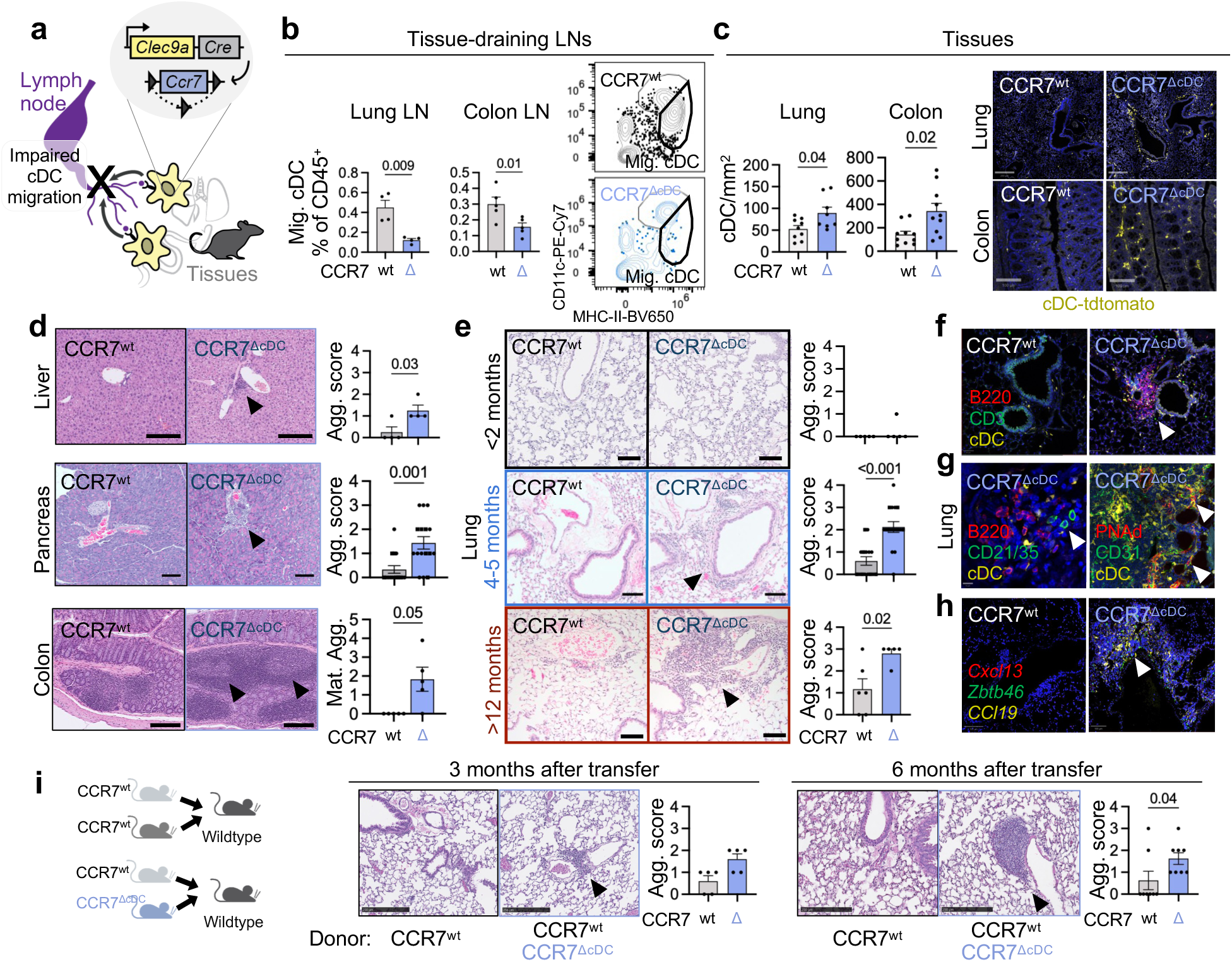
Tissue retention of cDCs is sufficient for de novo lymphoid organization in homeostatic tissues. a, Schematic of the *Clec9a*^Cre^;*Ccr7*^flox^ (CCR7^ΔcDC^) model, in which cDC migration from tissues to tissue-draining lymph nodes (LN) is impaired. **b,** Migratory cDCs among CD45+ cells in tissue-draining LNs from CCR7^wt^ (*n* = 4 lung, 5 colon) and CCR7^ΔcDC^ (*n* = 4 lung, 4 colon) mice, with representative gating. Two-tailed unpaired *t*-test; representative of two independent experiments. **c,** Representative immunofluorescence and quantification of tdTomato+ lineage-traced cDCs in peripheral tissues from *Clec9a*^Cre^;*Rosa26*^LSL-tdTomato^;CCR7^wt^ (*n* = 9 lung, 10 colon) and CCR7^ΔcDC^ (*n* = 8 lung, 10 colon) mice. Two-tailed unpaired *t*-test; data pooled from two independent experiments. Scale bars, 250 µm (lung) and 100 µm (colon). ***d,e***, Representative H&E images and lymphoid-aggregate or mature aggregates (colon) quantification in tissues from CCR7^wt^ (*n* = 4 liver, 15 pancreas, 5 colon, 15 lung) and CCR7^ΔcDC^ (*n* = 4 liver, 16 pancreas, 5 colon, 16 lung) mice at 3–4 months (**d**) or the indicated ages (<2 months: *n*= 5 CCR7^wt^, 5 CCR7^ΔcDC^; >12 months: 5 CCR7^wt^, 5 CCR7^ΔcDC^) (**e**). Two-tailed unpaired *t*-test; data pooled from two independent experiments. Scale bars, 200 µm (**d**) and 100 µm (**e**). **f,g,** Representative immunofluorescence of B220+ B cells, CD3+ T cells and tdTomato+ cDCs (**f**) and CD21/35+ follicular DCs, CD31+ PNAd+ high endothelial venules and tdTomato+ cDCs (**g**) within aggregates in lungs from CCR7^wt^ and CCR7^ΔcDC^ mice. **h,** In situ hybridization for *Cxcl13*, *Ccl19* and *Zbtb46* in lungs from CCR7^wt^ and CCR7^ΔcDC^ mice. **i,** Bone-marrow chimera design, representative H&E images and aggregate quantification 3 and 6 months after transfer (3 months: *n*= 5 CCR7^wt^, 5 CCR7^wt^+CCR7^ΔcDC^; 6 months: 8 CCR7^wt^, 8 CCR7^wt^+CCR7^ΔcDC^). Two-tailed unpaired *t*-test; data pooled from two independent experiments. Scale bars, 250 µm. Data are mean ± s.e.m.

We next asked whether cDCs had acquired a distinct transcriptional state as a consequence of CCR7 deficiency. Bulk RNA sequencing of cDCs isolated from spleen and lungs from young mice revealed minimal transcriptional divergence between CCR7^wt^ and CCR7^ΔcDC^ cells, with no differentially expressed genes at FDR < 0.05 and no distinct transcriptional program in either tissue (Extended Data Fig. 2h, Extended Data Table 1). Thus, CCR7-deficient cDCs did not acquire a substantial transcriptional program that distinguished them from control cDCs, supporting the idea that their tissue-organizing activity reflects their altered persistence and trafficking rather than the acquisition of a CCR7-dependent specialized phenotype.

Strikingly, in the absence of inflammatory challenge, CCR7^ΔcDC^ mice spontaneously developed de novo lymphoid aggregates in multiple peripheral organs, including liver, pancreas, salivary gland and lung, and showed an increased burden of mature lymphoid aggregates with germinal centers in the colon (Fig. 2d,e). Interestingly, lymphoid aggregate formation was age-dependent: structures were largely absent in young CCR7^ΔcDC^ mice at or before 2 months of age, started emerging in adult mice by 3–4 months and became progressively more prominent in older mice after 12 months of age (Fig. 2d,e and Extended Data Fig. 2i,h). Wild-type mice also developed age-associated lymphoid aggregates in tissues, but these arose later and were substantially less pronounced when compared to age-matched CCR7^ΔcDC^ (Fig. 1e, 2e, and Extended Data Fig. 1g, 2j). Importantly, in contrast to whole-body CCR7 deficient mice, which present systemic autoimmunity and disrupted thymic organization^25,26^, lymphoid aggregate-bearing CCR7^ΔcDC^ mice showed no histological tissue injury, no systemic inflammatory cytokine elevation in serum, no circulating autoantibodies, no renal immunoglobulin deposition, nor thymus disruption (Fig. 1d,e, Extended Data Fig. 2k-m, and Extended Data Table 2), indicating that lymphoid organization occurred in the absence of broad inflammatory or autoimmune pathology.

The resulting structures in mice with migration-deficient cDCs contained cDCs together with T and B lymphocytes (Fig. 2f and Extended Data Fig. 2n) and acquired canonical features of tertiary lymphoid structures (TLS), including the presence of CD21/CD35^+^ follicular DCs, PNAd^+^ high endothelial venules (Fig. 2g), and localized stromal expression of classical TLS-related cytokines *Cxcl13* and *Ccl19* (Fig. 2gh). Laser-capture microdissection-based spatial proteomics further revealed enrichment of pathways associated with functional TLS such as lymphocyte differentiation, leukocyte adhesion, immune activation and antigen presentation relative to normal tissue, indicating active immune stimulation within the aggregates (Extended Data Fig. 2n and Extended Data Tables 3,4). Immunoglobulin proteins were also detected within the aggregates, consistent with local B-cell differentiation and antibody production (Extended Data Table 3).

To physiologically model age-associated partial lack of cDC migration^21,22^ and exclude potential developmental defects due to the constitutive lack of CCR7 in cDCs, we generated mixed bone marrow chimeras. Adult wild-type recipients were reconstituted with either CCR7^wt^ bone marrow or a 1:1 mixture of CCR7^wt^ and CCR7^ΔcDC^ bone marrow, such that only a fraction of the cDC compartment was deficient for migration. Despite the presence of a substantial migration-competent cDC population, mixed chimeras developed age-dependent ectopic lymphoid aggregates in peripheral tissues, whereas control chimeras did not (Fig. 2i and Extended Data Fig. 2p). Thus, partial disruption of cDC egress is a dominant phenotype sufficient to trigger ectopic lymphoid organization in an otherwise normally developed, migration-competent immune system.

Collectively, these findings establish that a partial defect in mature cDCs migration from tissues to draining LNs is a sufficient driver of age-dependent de novo lymphoid organization in otherwise homeostatic tissues, generating structures with canonical features of tertiary lymphoid niches in absence of associated tissue injury.

### Senescence drives lymphoid aggregates

Because migratory cDCs play an important role for peripheral tolerance in tissue-draining LNs, we next asked whether established lymphoid structures remained dependent on the accumulation of cDC in the tissue over an absence of migratory cDCs reaching the draining LN. Acute depletion of cDCs from TLS-bearing adult CCR7^ΔcDC^;*Zbtb46*^DTR^ mice markedly dissolved established aggregates within 1 week of diphtheria toxin (DT) treatment. This demonstrates that established age-associated TLS require continuous presence of migratory cDC accumulation in tissues (Fig. 3a and Extended Data 3a,b).

**Fig. 3.**
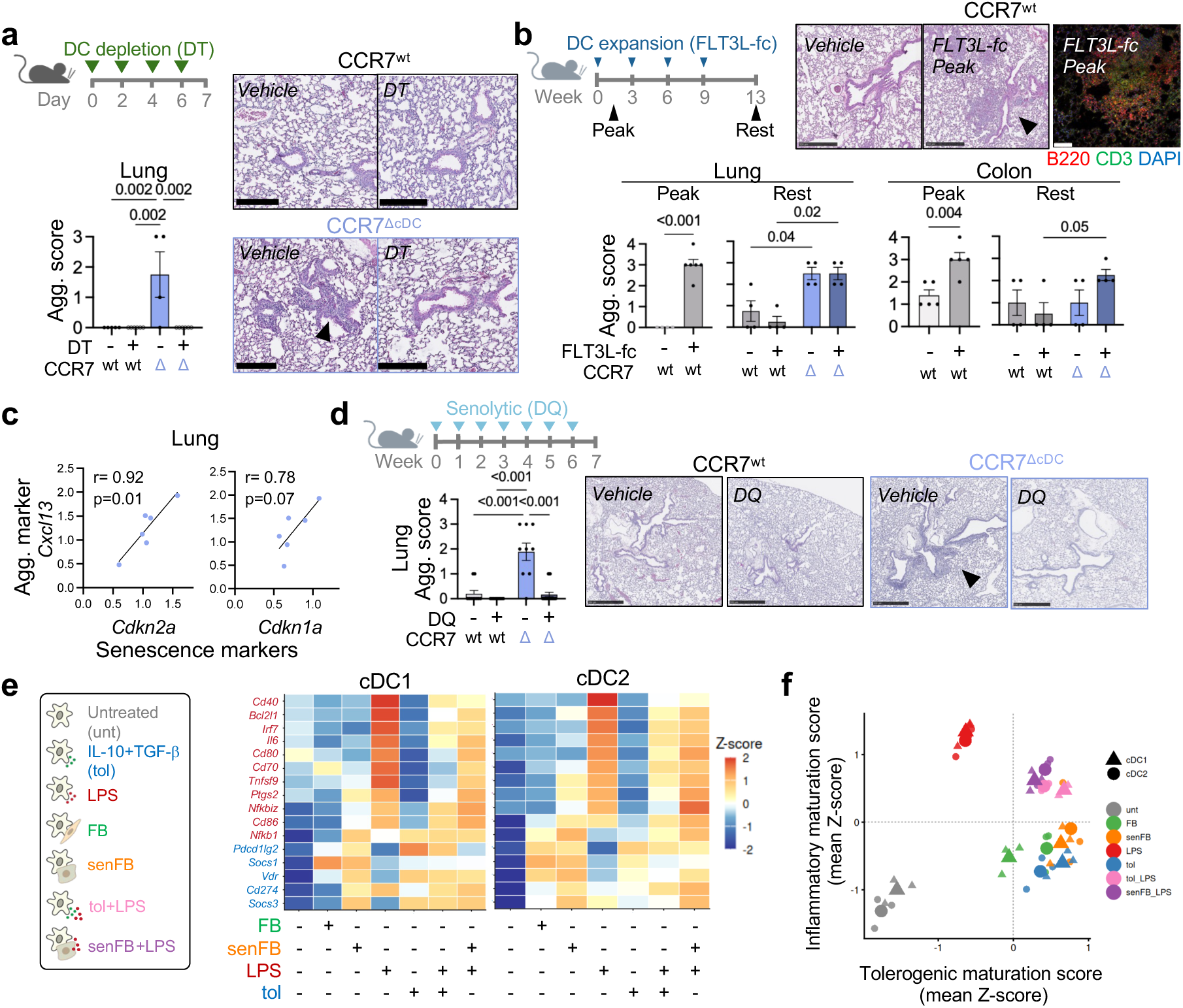
Age-associated senescence shapes cDC maturation state to organize lymphoid niches in tissues. a, Experimental design, representative H&E images and aggregate quantification in CCR7^wt^;*Zbtb46*^DTR^ (*n* = 5 untreated, 6 treated) and CCR7^ΔcDC^;*Zbtb46*^DTR^ (*n* = 4 untreated, 6 treated) adult mice untreated or after 1 week of diphtheria toxin-mediated cDC depletion. One-way ANOVA with two-stage step-up correction; representative of two independent experiments. Scale bars, 250 µm. **b,** Experimental design, representative H&E and immunofluorescence images and aggregate quantification in wild-type adult mice following FLT3L–Fc-induced cDC expansion at peak expansion (day 12) (*n* = 4 untreated lung 5 untreated colon; 6 treated lung, 5 treated colon) and 30 days after the final treatment (*n* = 4 untreated CCR7^wt^, 4 treated CCR7^wt^, 4 untreated CCR7^ΔcDC^, 4 treated CCR7^ΔcDC^). Immunofluorescence shows B220+ B cells, CD3+ T cells and DAPI (*n* = 4–6). Two-tailed *t*-test and one-way ANOVA with two-stage step-up correction. Scale bars, 500 µm (H&E) and 100 µm (immunofluorescence). **c,** Correlations between qPCR-normalized lung expression of the senescence markers *Cdkn2a* or *Cdkn1a* and the aggregate marker *Cxcl13* in adult CCR7^ΔcDC^ mice (*n* = 6). Linear regression with Pearson *r* and two-tailed *P* values. **d,** Experimental design, representative H&E images and lymphoid-aggregate quantification in young 2-month-old CCR7^wt^ (*n* = 10 untreated, 12 treated) and CCR7^ΔcDC^ (*n* = 9 untreated, 15 treated) mice treated for 7 weeks with vehicle or dasatinib plus quercetin (DQ). One-way ANOVA with two-stage step-up correction. Data pooled from two experiments. Scale bars, 500 µm. **e,** Experimental design and heatmap of mean z-scored log2 counts per million (CPM) for selected cDC inflammatory(red) and tolerogenic maturation (blue) signature genes in cDC1 (*n* = 3) and cDC2 (*n* = 3) across culture conditions. FB, fibroblast; senFB, senescent fibroblast; LPS, lipopolysaccharide; tol, tolerogenic factors. **f,** Projection of cDCs across same culture conditions showing immunogenic and tolerogenic signature scores calculated from mean z-scored log2-CPM values. Data are mean ± s.e.m.

To determine whether cDC-driven lymphoid-organizing activity could be pharmacologically triggered by skewing the hematopoiesis toward the myeloid lineage with FLT3L growth factor to increase cDC abundance in tissues, we expanded cDCs in wild-type adult mice using FLT3L-Fc^27^. At the peak of cDC expansion, wild-type adult mice developed ectopic lymphoid aggregates in tissues such as lung and salivary gland and showed increased aggregate burden and maturation in the colon (Fig. 3b and Extended Data Fig. 3c,d). These structures contained both B and T lymphocytes (Fig. 3d and Extended Data Fig. 3d), demonstrating that increased cDC abundance in tissue is sufficient to initiate ectopic lymphoid aggregate organization in otherwise homeostatic organs. We next asked whether these niches persisted once cDC burden returned towards baseline. Following 3 months of chronic FLT3L-Fc treatment and 1-month rest period after withdrawal, induced aggregates largely regressed in wild-type mice, whereas aggregates in CCR7^ΔcDC^ mice persisted and were further increased relative to untreated controls in several tissues (Fig. 3b and Extended Data Fig. 3e). These findings distinguish between initiation from maintenance of lymphoid aggregates: transient cDC expansion is sufficient to trigger lymphoid organization, whereas sustained tissue persistence of migratory cDCs is required for long-term maintenance of the aggregates in mice.

The age dependence of lymphoid aggregate formation in CCR7^ΔcDC^ mice suggested that cDC persistence cooperates with tissue changes that emerge with aging. We therefore asked whether cellular senescence, a hallmark of aging^14^ contributes to cDC-driven lymphoid organization. Indeed, in aggregate-bearing lungs from CCR7^ΔcDC^ mice, expression of the senescence-associated genes *Cdkn1a* and *Cdkn2a* correlated with expression of the TLS-associated chemokine *Cxcl13* (Fig. 3c).

To test whether senescence-associated signals contribute to niche initiation, we treated young CCR7^ΔcDC^ mice before aggregate formation with the senolytic combination dasatinib and quercetin (DQ), which specifically targets senescent cells by disrupting their pro-survival pathways^25^ (Fig. 3d). DQ treatment reduced the expression of the senescence-associated markers *Cdkn1a* and *Cdkn2a* in tissues, without altering systemic abundance of cDCs (Extended Data Fig. 3f,g). Remarkably, senolytic treatment markedly prevented lymphoid aggregate formation in CCR7^ΔcDC^ mice (Fig. 3d, Extended Data 3h). By contrast, the same treatment did not dissolve pre-existing structures when administered to adult CCR7^ΔcDC^ mice with already established aggregates in tissues (Extended Data Fig. 3i). Thus, senescence-associated tissue cues are required for the initiation of cDC-driven lymphoid niches but become dispensable once the structures are established.

To study the effects of age-associated tissue senescence on cDCs state, we co-cultured primary mouse cDCs (both cDC type 1 and 2) isolated from spleens of FLT3L-Fc-treated mice with primary fibroblasts rendered senescent by replicative exhaustion (Extended Data Fig. 3j) and analysed cDC state by bulk RNAseq. We benchmarked the resulting transcriptional state against inflammatory (LPS) or tolerogenic (IL-10 plus TGF-β) cDC maturation conditions (Fig. 3e). Unbiased principal-component (PC) analysis showed that cDCs exposed to senescent fibroblasts adopted a transcriptional state distinct from unstimulated cDCs and from both inflammatory and tolerogenic factors (Extended Data Fig. 3k). Interestingly, senescence-exposed cDCs were transcriptionally closer to cDCs treated with tolerogenic factors (Extended Data Fig. 3k). We found that senescent fibroblasts induced a unique maturation program with limited upregulation of certain maturation markers induced by inflammatory maturation and a wide upregulation of many genes induced by tolerogenic factors (Figure 3e and Extended Data Fig. 3l, Extended Data Table 5). Consistent with this, projection onto inflammatory and tolerogenic maturation gene signatures placed senescence-exposed cDCs in an intermediate state characterized by a mild upregulation of inflammatory maturation together with a strong tolerogenic maturation signature (Figure 3f). Moreover, coculture with senescent fibroblasts dampened the inflammatory transcriptional maturation program driven by LPS in cDCs exposed to both stimuli with similar strength than tolerogenic factors (Figure 3e,f and Extended Data Fig. 3k,l).

Taken together, these results suggest that age-associated tissue cues promote a non-classical homeostatic cDC maturation with regulatory features, while increased abundance and sustained persistence of those cDCs in tissues provide the organizing force required to initiate and maintain lymphoid niches.

### cDC niches enrich tissue-adapted Tregs

To assess the resulting function of cDC-driven age-associated lymphoid niches on adaptive immunity, we then examined the lymphocyte composition of aggregate-bearing tissues using flow cytometry. Unexpectedly, the dominant relative change across tissues from CCR7^ΔcDC^ mice was the enrichment of FOXP3^+^ regulatory T cells (Tregs) (Fig. 4a,b and Extended Fig. 4a,b). Immunofluorescence analysis revealed that FOXP3^+^ Tregs preferentially accumulated within lymphoid aggregates rather than throughout the surrounding tissue parenchyma (Fig. 4c). Within lymphoid niches, the proportion of FOXP3^+^ cells among total T cells was increased in CCR7^ΔcDC^ mice, indicating a selective enrichment of Treg rather than a proportional increase in all T-cell subsets. In the colon, where lymphoid aggregates are also present in wild-type mice, both the abundance of Treg and the proportion of Tregs out of total T cells within lymphoid aggregates were increased in CCR7^ΔcDC^ mice (Fig. 4c).

**Fig. 4.**
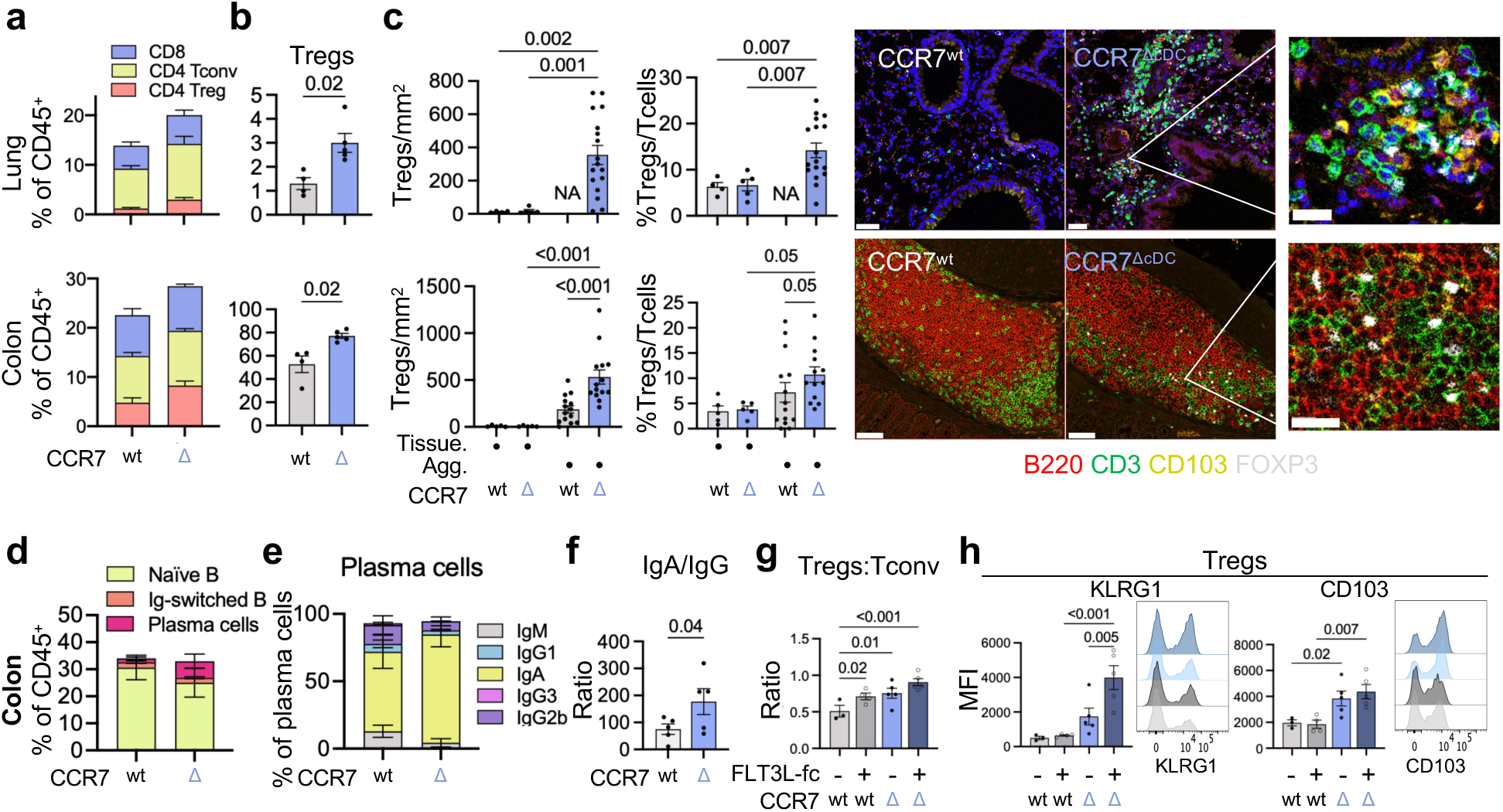
cDC-driven lymphoid structures are enriched in regulatory features. a,b, Flow-cytometric quantification of CD8+ T cells, CD4+ conventional T (Tconv) cells and FOXP3+ regulatory T (Treg) cells (**a**) and Treg cells as a percentage of CD45+ cells (**b**) in peripheral tissues from CCR7^wt^ (*n* = 4 lung, 4 colon) and CCR7^ΔcDC^ (*n* = 5 lung, 5 colon) mice. Two-tailed unpaired *t*-test in **b**; representative of two independent experiments. **c,** FOXP3+ Treg-cell density, Treg cells as a proportion of CD3+ T cells and representative immunofluorescence of B220+ B cells, CD3+ T cells, CD103+ cDCs and FOXP3+ Treg cells within lymphoid aggregates. Each dot represents random tissue areas averaged per mouse or an independent lymphoid aggregate from CCR7^WT^ (*n* = 4 lung, 5 colon) and CCR7^ΔcDC^ (*n* = 5 lung, 5 colon) mice. One-way ANOVA with two-stage step-up correction. Scale bars, 50 µm and 20 µm (zoomed images). **d,e,** Flow-cytometric quantification of naive B cells, IgD−IgM− class-switched B cells and plasma cells (**d**) and plasma-cell immunoglobulin isotypes (**e**) in colon from CCR7^WT^ and CCR7^ΔcDC^ mice (*n* = 5–6). **f,** Fecal IgA/IgG ratio in feces from CCR7^WT^ (*n* = 5) and CCR7^ΔcDC^ (*n* = 5) mice. Two-tailed unpaired *t*-test. **g,** Treg:Tconv ratios following FLT3L–Fc-induced cDC expansion in colon from CCR7^WT^ (*n* = 3 untreated, 4 treated) and CCR7^ΔcDC^ (*n* = 5 untreated, 5 treated) mice. **h,** KLRG1 and CD103 expression by colonic Treg cells from the same groups. One-way ANOVA with two-stage step-up correction in **g,h**. Data are mean ± s.e.m.

Because the colon of CCR7^ΔcDC^ mice contained the most mature lymphoid structures, including germinal-center-containing aggregates (Fig. 2d), we further examined B-cell differentiation in this tissue. This Treg-enriched regulatory composition was accompanied by increased non-naive B cell and plasma-cell differentiation in the colon and an IgA-skewed mucosal antibody response in feces (Fig. 4d-f, Extended Data Fig. 4d). Together, these findings indicate that cDC-driven lymphoid niches are biased toward regulatory mucosal immunity, characterized by Treg enrichment and an IgA-skewed B-cell response in the colon tissue.

To determine whether pharmacological increase of cDC abundance was sufficient to promote this regulatory bias, we expanded cDCs with FLT3L-Fc treatment. In wild-type mice, cDC expansion preferentially increased Treg abundance and elevated Treg-to-conventional CD4 and Treg-to-CD8 T cell ratios to levels approaching those observed in CCR7^ΔcDC^ mice in multiple tissues (Fig. 4g and Extended Data Fig. 4e). In CCR7^ΔcDC^ mice, cDC expansion with FLT3L-Fc further enhanced those regulatory phenotypes (Fig. 4g and Extended Data Fig. 4d,e). Conversely, the acute cDC depletion in CCR7^ΔcDC^;*Zbtb46*^DTR^ mice reduced Treg abundance and the Treg-to-conventional T-cell ratio towards wild-type levels (Extended Data Fig. 4f). Thus, continued mature cDC accumulation in tissue is required to promote and to sustain the Treg-biased composition of these lymphoid niches.

Phenotypic profiling showed that Tregs from CCR7^ΔcDC^ tissues expressed increased levels of markers associated with tissue residency and effector Treg differentiation, including KLRG1, CD103, ST2 and CD49b (Fig. 4h and Extended Data Fig. 4g). These findings indicate that cDC-driven niches preferentially accumulate a tissue-adapted effector Treg population rather than simply increasing total Treg numbers.

Overall, these findings show that cDC-driven age-associated lymphoid niches preferentially accumulate tissue-adapted Tregs and are associated with a regulatory mucosal immune phenotype.

### cDC retention limits tissue injury

We next tested whether the immune environment created by persistent tissue cDCs and associated lymphoid niches have broader functional consequences for tissue biology. For that, we challenged CCR7^ΔcDC^ mice with DSS-induced colitis, as a model of chemically-induced colon tissue injury and repair. Compared with CCR7^wt^ controls, CCR7^ΔcDC^ mice showed improved recovery from one cycle of DSS-induced damage, accompanied by reduced colon shortening and mucosal inflammation and ulceration (Fig. 5a,b). Differences were not apparent during the initial phase of epithelial injury but emerged after DSS withdrawal, beginning around days 8–10 during tissue recovery. This delayed protection therefore suggested an effect on immune-mediated resolution and repair rather than reduced susceptibility to the initial barrier insult. We next extended these findings to a chronic DSS model, in which repeated cycles of epithelial injury generate sustained inflammation and greater involvement of adaptive immunity. Under these conditions, differences became increasingly evident over time, indicating that the protective phenotype was sustained even during recurrent tissue injury (Extended Data Fig. 5a). Similar protection against chronic damage was also observed in mixed bone-marrow chimeric mice with a partial defect of cDC migration within an otherwise migration-competent haematopoietic compartment (Extended Data Fig. 5b), supporting a dominant tissue cDC-dependent protective effect rather than a consequence of constitutive CCR7 deficiency in cDCs.

**Fig. 5.**
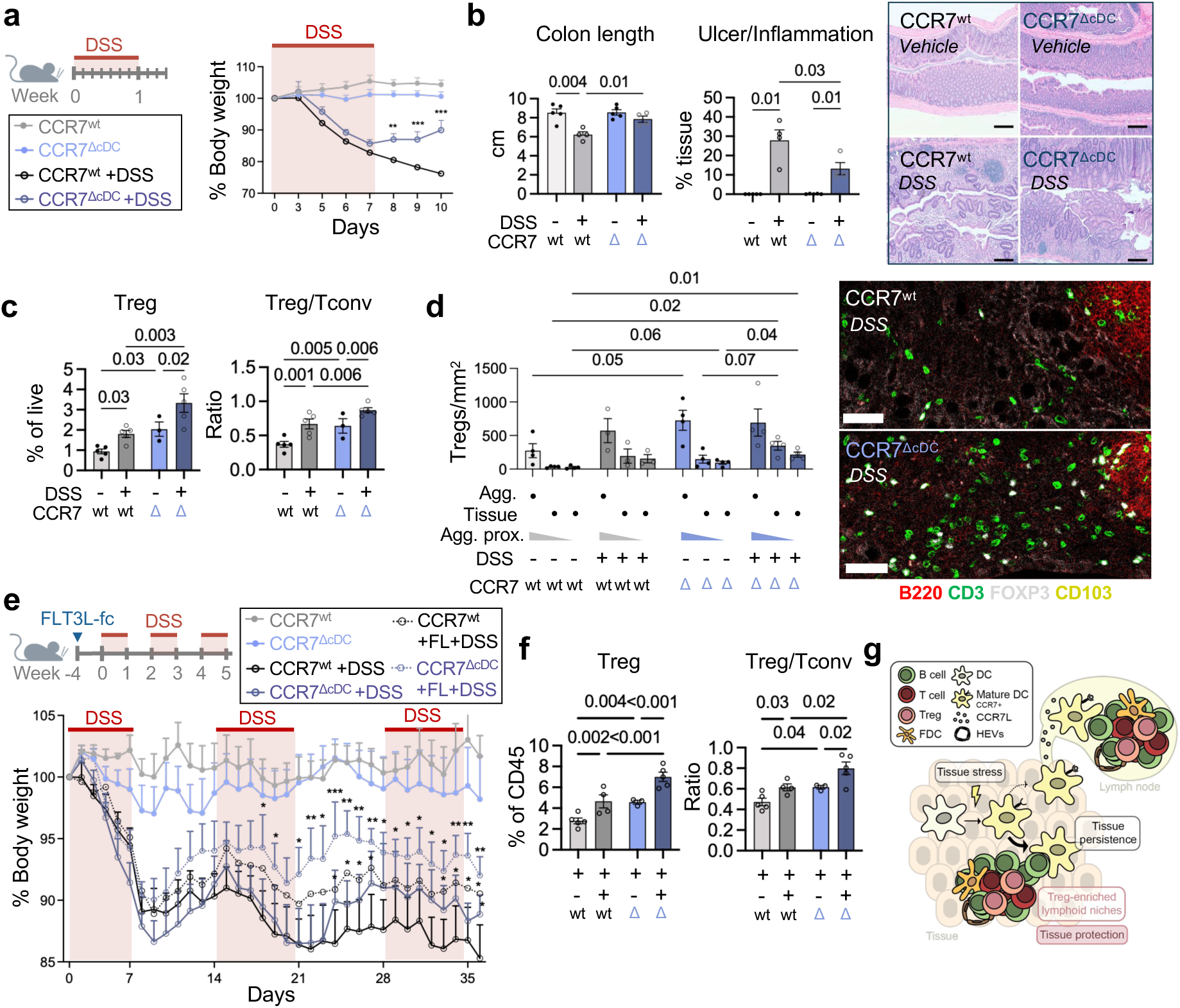
cDC persistence in tissues limits inflammatory injury. a, Acute colitis induced by 3% dextran sodium sulfate (DSS) for 7 days and longitudinal body weight in CCR7^WT^ (*n* = 5 vehicle, 4 DSS) and CCR7^ΔcDC^ (*n* = 5 vehicle, 4 DSS) mice. Two-way ANOVA with two-stage step-up correction; representative of two independent experiments. Indicated significant p values are against DSS-treated CCR7^WT^ mice. **b,** Colon length, ulcerated or inflamed mucosal area and representative H&E images from untreated and DSS-treated CCR7^WT^ (*n* = 5 vehicle, 4 DSS) and CCR7^ΔcDC^ mice (*n* = 5 vehicle, 4 DSS). One-way ANOVA with two-stage step-up correction. Scale bars, 200 µm. **c,** Flow-cytometric analysis of Treg-cell abundance and Treg ratio in colon from CCR7^WT^ (*n* = 5 vehicle, 5 DSS) and CCR7^ΔcDC^ mice (*n* = 3 vehicle, 5 DSS) at day 7 after DSS. One-way ANOVA with two-stage step-up correction; representative of two independent experiments. **d,** Representative immunofluorescence and quantification of FOXP3+ Treg cells within lymphoid aggregates and adjacent or distal tissue in colon from CCR7^WT^ (*n* = 4 vehicle, 3 DSS) and CCR7^ΔcDC^ mice (*n* = 4 vehicle, 4 DSS) at day 7 after DSS. Each dot represents random tissue areas averaged and lymphoid aggregates per mouse. Two-way ANOVA with Fisher’s LSD test. Scale bars, 50 µm. **e,** Chronic colitis induced by three 7-day cycles of 2% DSS separated by 7-day recovery periods, with or without FLT3L–Fc pretreatment, and longitudinal body weight CCR7^WT^ (*n* = 5 untreated, *n* = 6 FLT3L-fc-treated DSS, 6 treated DSS) and CCR7^ΔcDC^ mice (*n* = 5 untreated, 5 FLT3L-treated DSS, 4 untreated DSS). Two-way ANOVA with multiple comparisons; indicated significant p values are against DSS-treated CCR7^WT^ mice. **f,** Flow-cytometric analysis of Treg-cell abundance and Treg ratio from CCR7^WT^ (*n* = 5 untreated DSS, 4 treated DSS) and CCR7^ΔcDC^ (*n* = 3 untreated DSS, 5 treated DSS) mice at day 7 after DSS at day 7 after DSS with or without FLT3L–Fc treatment. Two-way ANOVA with two-stage step-up correction. **g,** Working model of tissue-stress-induced cDC maturation and cDC persistence driving Treg-rich lymphoid-aggregate formation and tissue protection. Data are mean ± s.e.m.

To characterize the immune state associated with tissue protection, we analysed using flow cytometry the colonic immune compartment at day 7 after DSS administration, when both CCR7^wt^ and CCR7^ΔcDC^ mice displayed comparable weight loss, allowing us to identify potential immune differences preceding the emergence of protection (Fig. 5a). Colons from CCR7^ΔcDC^ mice contained a similar burden of T conventional CD4 T cells (Tconv) but an increased Tregs abundance and an increased Treg-to-conventional T-cell ratio in comparison to CCR7^wt^ mice (Fig. 5c and Extended Data Fig 5c,d). To further explore the spatial distribution of Tregs in colon tissue after DSS-induced damage, we perform immunofluorescence analysis. Similarly to uninjured tissue, Tregs preferentially accumulated within organized lymphoid aggregates in damaged tissue, but were also increased in the surrounding lamina propria suggesting a relocation of tissue-adapted Tregs in response to damage (Fig. 5d). Moreover, DSS-treated CCR7^ΔcDC^ mice contained more Tregs in the lamina propria than DSS-treated CCR7^wt^ mice (Fig. 5d). The proportion of Tregs among total T cells was elevated across tissue compartments in CCR7^ΔcDC^ mice, indicating selective Treg enrichment rather than generalized T-cell accumulation (Extended Data Fig. 5e). These observations suggested that cDC tissue persistence and associated lymphoid niches support a preferential accumulation of Tregs in injured tissues.

We next asked whether the protective state induced by persistent cDC accumulation and associated lymphoid aggregates in tissues could be induced by increasing cDC abundance as a therapeutic intervention. FLT3L-Fc treatment before chronic DSS challenge attenuated weight loss in CCR7^wt^ mice, demonstrating that increasing cDC abundance alone is sufficient to promote tissue protection (Fig. 5e). Notably, protection was further enhanced in CCR7^ΔcDC^ mice, suggesting that cDC expansion and tissue retention act synergistically to increase tissue resilience. Consistent with this protective phenotype, FLT3L-Fc-mediated cDC expansion before DSS-induced damage increased the proportion of colonic Tregs and the Treg-to-conventional T-cell ratio in both genotypes at day 7 after DSS administration, without increasing Tconv burden (Fig. 5f and Extended Data Fig. 5f). Thus, pharmacological expansion of cDCs can recapitulate key features of the protective immune state associated with cDC persistence, linking increased cDC abundance to enhanced regulatory T-cell representation during tissue injury.

In conclusion, our observations identify cDC accumulation in tissues as a feature of a regulatory, tissue-protective immune state and link cDC-driven lymphoid organization with enhanced tissue resilience during inflammatory tissue injury that could be therapeutically leveraged for chronic inflammatory diseases (Fig. 5g).

## Discussion

Our study establishes that ectopic lymphoid aggregates, including tertiary lymphoid structures (TLS), are not necessarily inflammatory, as often viewed in the context of cancer, infection or autoimmune disease^1,4^. Instead, they can also represent a form of lymphoid organization that arises spontaneously across tissues with age and is enriched in regulatory features. These structures may therefore represent an organized immune adaptation arising to promote tissue tolerance and protection in aged tissues in which naturally-occurring senescence and impaired immune-cell egress progressively accumulate^14,21,22^. This may be especially relevant at mucosal interfaces such as the intestine, where immune activation must remain tightly coupled with tissue tolerance and barrier preservation.

Our work further identifies a novel paradigm of tissue retention as an important determinant of conventional dendritic cell (cDC) function. cDCs maturation is normally coupled to CCR7-dependent migration from peripheral tissues to tissue-draining lymph nodes (LNs), where cDC coordinate adaptive immunity and tolerance. Recently, cDCs have been shown to regulate the formation and maintenance of cancer-associated inflammatory TLS, a process requiring both cDC maturation and CCR7-dependent migration to tumor-draining LNs^28^. Here, we decouple cDC migration from maturation and show, at the opposite, that selective impairment of cDC egress from tissues to LNs accelerated the formation of age-associated regulatory lymphoid niches with canonical features of TLS across multiple otherwise homeostatic peripheral organs. Together, these findings suggest that the signals and cellular context initiating ectopic lymphoid organization may be important determinants of whether the resulting niche supports effector or regulatory immunity.

Finally, this framework raises the possibility that cDC localization, abundance and persistence could be manipulated to reshape tissue immune states. Both pharmacological expansion of the cDC compartment and restriction of cDC egress from tissues, alone and especially in combination, induced ectopic lymphoid organization and features of regulatory tissue protection. Thus, therapeutic modulation of cDCs in tissues may provide a potential opportunity to shift local immune niches toward regulatory rather than inflammatory states in chronic inflammatory diseases, autoimmunity or organ transplantation.

## Methods

### Human tissue

Human tissues were acquired from multiple vendor sources including Avaden Biosciences, Analytical Biological Services, Bio-options Inc, Bioreclaimation IVT, Clinomics Laboratories, ClinPath Advisors, Cooperative Human Tissue Network, Cureline, FFPExpress, Folio, Indivumed, Precision Biospecimen Solutions, and the University of Michigan. Providers ensured necessary informed consent for research use.

### Mice and in vivo treatments

*Clec9a^Cre^CCR7^fl/fl^* (CCR7^ΔcDC^), *Clec9a^Cre^CCR7^fl/fl^Rosa26^LSL.tdtomato^*, and *Clec9a^Cre^CCR7^fl/fl^Zbtb46^DTR^* (CCR7^ΔcDC^Zbtb46^DTR^), mice were designed, generated and bred at Genentech. For mouse experiments, littermate Cre-expressing CCR7 wild-type mice (CCR7^wt^) were used as controls. Male and female adult mice aged 4-5 months were used for all studies unless specified. C57BL/6 (Jax: 000664) and CD45.1 C57BL/6 (Jax: 002014) mice were obtained from Jackson Laboratories Inc. (Hollister, CA). Mice were housed in specific-pathogen-free, individually ventilated cages following the guidelines of the US National Institutes of Health. Animal rooms were maintained on a 14:10 h light–dark cycle and were temperature-controlled and humidity-controlled at 20–26 °C and 30–70% humidity, respectively, with 10–15 room air exchanges per hour. For cDC depletion in CCR7^ΔcDC^Zbtb46^DTR^ mice, mice were intraperitoneally injected with 25 ng g−1 of diphtheria toxin (DT) (Enzo Life Sciences, BML-G135) every other day. For cDC expansion, mouse FLT3L-Fc was administered at 2mg/kg intraperitoneally at the indicated timepoints. For senolytic treatment, mice were intraperitoneally injected with the senolytic cocktail 10mg/kg dasatinib (Sigma-Aldrich, CDS023389) and 50mg/kg quercetin (Sigma-Aldrich, Q4951) on a weekly basis. All mouse studies were approved by the Genentech Institutional Animal Care and Use Committee or the Animal Care Committee at TCP. No statistical methods were used to predetermine sample size. The experiments were randomized. The investigators were not blinded to allocation for the experiments, but they were for outcome assessment.

### Serum preparation from mouse blood

Whole blood was collected from mice into serum-separation tubes without anticoagulants. Samples were allowed to clot at room temperature for 30 min and centrifuged at 2,000 × g for 10 min at 4 °C. The clear serum fraction was transferred to fresh tubes without disturbing the cell pellet, aliquoted and stored at −80 °C until analysis.

### Anti-nuclear antibodies

Antinuclear antibodies in mouse serum were detected by indirect immunofluorescence using HEp-2 cell substrate slides (Biorad, 26102) following manufacturer instructions. Briefly, serum samples were diluted 1:50 in PBS and incubated on the slides for 30 min at room temperature in a humidified chamber. Slides were washed three times with PBS and incubated for 30 min with an FITC-conjugated anti-mouse IgG secondary antibody (1:500). After three further washes, slides were mounted with ProLong Gold Antifade mounting medium (Invitrogen) and imaged using a fluorescence microscope. Positivity and staining patterns were assessed relative to positive and negative control sera.

### Ear FITC painting migration assay

Mice were anesthetized with isoflurane. The dorsal and ventral surfaces of the right ear were painted with 0.5% FITC solution (prepared in a 1:1 mixture of acetone and dibutyl phthalate) and allowed to air dry for 1–10 minutes under anesthesia. Ear-draining LNs were analysed 48 hours later.

### Bone marrow chimera generation

Bone marrow chimeras were generated by lethally irradiating recipient 8–12-week-old CD45.1 C57BL/6 (Jax: 002014) female mice with 10.5 Gy total body irradiation delivered as a split dose (5 Gy and 5.5 Gy), administered 3 hours apart. Recipient mice were reconstituted intravenously via the retro-orbital plexus with up to 20 million bone marrow cells harvested from donor mice CD45.1 C57BL/6 (Jax: 002014) or CD45.2 CCR7^ΔcDC^ or CCR7^ΔcDC^ mice, as specified. Donor bone marrow was prepared by flushing femurs and tibias with ice-cold PBS, red blood lysis with ACK, and passing through a 70 μm cell strainer. Hematopoietic reconstitution was verified at the end of the experiment by flow cytometry analysis of spleen (CD45.1/CD45.2 congenic markers). Bone marrow chimeric mice were used 12 weeks post-transplant to allow complete hematopoietic reconstitution for DSS-induced colitis or aged for subsequent analysis of tissue aggregate formation.

### DSS-induced colitis model

Acute colitis was induced by administering dextran sulfate sodium (DSS; molecular weight 36–50 kDa; MP Biomedicals) in drinking water at a concentration of 3% for 7 days followed by normal water. Control mice received normal drinking water. Chronic colitis was induced using a repeated DSS protocol. Mice received 2% or 2.5% DSS (as indicated in the figure legend) in drinking water for 7 days, followed by 7 days of normal water. This cycle was repeated up to 3 times over 6 weeks. Body weight was monitored throughout the treatment period. Control mice received normal drinking water. In both acute and chronic colitis, body weight was monitored as a readout of disease throughout the treatment period. Disease score was assessed evaluating stool consistency, occult blood, and gross blood in feces on a scale of 0–4. Mice were euthanized at the indicated timepoint, and colons were isolated, measured, and processed for downstream analysis.

### Leukocyte isolation from mouse tissues

Colons were cleaned of fat, flushed with PBS and incubated in prewarmed IEL buffer consisting of PBS, 2% FBS, 10 mM HEPES, 1% penicillin–streptomycin, 1% L-glutamine or GlutaMAX, 5 mM EDTA and 1 mM dithiothreitol for 15 min at 37 °C with shaking at 250 r.p.m. Samples were washed with PBS and digested in prewarmed LPL buffer consisting of RPMI 1640, 2% FBS, 10 mM HEPES, 1% penicillin–streptomycin, 1% L-glutamine or GlutaMAX, 1 mg ml−1 collagenase A and 1 U ml−1 DNase I for 30 min at 37 °C with shaking at 250 r.p.m. Samples were filtered through a 100-µm strainer, centrifuged and washed using cold MACS buffer. When necessary, immune cells were further cleaned up with Percoll 40%.

Lungs were mechanically minced into small pieces and digested in RPMI containing 2% FBS, 0.5 mg ml−1 collagenase IV and 0.25 mg ml−1 DNase I for 30 min at 37 °C with shaking at 900 r.p.m. Digested tissue was passed through a 100-µm filter pre-wetted with MACS buffer and washed with ice-cold MACS buffer. Cells were centrifuged, resuspended in ACK lysis buffer and incubated for 5 min. Lysis was stopped with ice-cold MACS buffer. Samples were filtered through a 70-µm strainer, centrifuged, and resuspended in MACS buffer.

Spleens and lymph nodes were mechanically minced into small pieces and digested in RPMI containing 2% FBS, 1 mg ml−1 collagenase D and 0.1 mg ml−1 DNase I for 25 and 20 min, respectively, at 37 °C with shaking at 900 r.p.m. Digested tissue was passed through a 70-µm filter pre-wetted with MACS buffer and washed ice-cold MACS buffer. Cells were centrifuged, resuspended in ACK lysis buffer and incubated for 5 min. Lysis was stopped using a cold MACS buffer.

### Flow cytometry analysis

Cells were then Fc blocked with anti-CD16/32 (BD Pharmigen) for 10 minutes on ice, followed by incubation with fluorescently labeled surface antibodies for 25 minutes on ice. After surface staining, cells were fixed and permeabilized using a Foxp3/transcription factor fixation/permeabilization staining buffer set (eBioscience) following manufacturer instructions, then stained with fluorescently labeled intracellular antibodies for at least 2h on ice. Cells analyzed on a LSRFortessa, BD FACSymphony, or Cytek Aurora flow cytometer and using FlowJo (v.10.8.1 or v.10.9.0) and Prism (v11). All antibodies are listed in Extended Data Table 6.

### cDC and senescent cell coculture

Primary mouse fibroblasts were isolated from ear tissue. Briefly, ears were mechanically minced and enzymatically digested with Dispase/Collagenase (Roche) and DNase I (Roche) for 30 min at 37°C. The digested tissue fragments were then cultured in complete DMEM media, allowing fibroblasts to migrate out of the tissue and expand in culture. Replicative senescence was induced by serial passaging of primary fibroblasts until proliferative arrest was reached, cells acquired a characteristic senescent morphology, and the majority of cells became positive for senescence-associated β-galactosidase (SA-β-gal) staining^29^.

Mouse primary DC were obtained from the spleens of FLT3L-fc treated mice. DC were enriched using the Pan-DC enrichment kit (Miltenyi). DC were cultured for 18 h in complete RPMI media alone, containing LPS (50 ng/mL), mouse recombinant IL-10 (10 ng/mL) and mouse recombinant TGF-beta 1 (10 ng/mL), primary mouse fibroblasts or senescent mouse fibroblasts.

### RNA extraction

Tissues were disrupted using a mechanical tissue processor in beads-containing tubes in RLT lysis buffer (Qiagen). Cells were directly resuspended by pipetting in RLT lysis buffer (Qiagen). RNA was extracted from tissues using the RNAeasy Mini Kit (Qiagen).

### Reverse transcription and quantitative PCR

Total RNA (up to 2 µg per sample) was reverse-transcribed using the High-Capacity cDNA Reverse Transcription Kit (Applied Biosystems) with random primers, according to the manufacturer’s instructions. Quantitative PCR was performed using TaqMan Fast Advanced Master Mix and gene-specific TaqMan probes *Cdkn2a* (Mm00494449_m1), *Cdkn1a* (Mm00432448_m1), *Cxcl13* (Mm00444533_m1), *Gapdh* (Mm99999915_g1) and *Actb* (Mm02619580_g1) (Thermo Fisher Scientific) following manufacturer instructions. Each sample was analyzed in technical triplicate. Relative gene expression was calculated after normalization to the averaged expression of the reference genes Actb and Gapdh .

### Bulk RNA sequencing of sorted cDC

cDC (Live CD45⁺Ly6C⁻F4/80⁻CD19⁻CD3⁻CD11c⁺MHCII⁺) were sorted from the lung and spleen of CCR7^ΔcDC^ and CCR7^wt^ littermate control mice (*n* = 4 per group per tissue). RNA was extracted as explained above. RNA was quantified with the Qubit RNA HS Assay Kit (Thermo Fisher Scientific) and quality was assessed using RNA ScreenTape on TapeStation 4200 (Agilent Technologies). cDNA library was generated from 2 nanograms of total RNA using the Smart-Seq V4 Ultra Low Input RNA Kit (Takara). 150 picograms of cDNA were used to make sequencing libraries with the Nextera XT DNA Sample Preparation Kit (Illumina). Libraries were quantified with the Qubit dsDNA HS Assay Kit (Thermo Fisher Scientific) and the average library size was determined using D1000 ScreenTape on TapeStation 4200 (Agilent Technologies). Libraries were pooled and sequenced on NovaSeq X Plus (Illumina) to generate 30 million single-end 50-base pair reads for each sample.

### Bulk RNAseq data processing of sorted mouse tissue cDC

Strand-unspecific RNA-seq libraries were prepared and sequenced at the Genentech Sequencing facility as explained above. Raw sequencing reads were processed through the Roche internal Sunrise platform using the congee_cromwell_prd-ngs_pipeline workflow. Raw read counts were analysed in R v4.6. Differential expression (DE) was performed separately for each tissue using the limma/voom pipeline (limma v3.x, edgeR v3.x)^30,31^. Briefly, genes with insufficient counts were removed using filterByExpr with default parameters, and library sizes were normalised using the trimmed mean of M-values (TMM) method. Precision weights were estimated with voomWithQualityWeights to account for sample-level heteroscedasticity^32^. A linear model was fitted with lmFit and empirical Bayes shrinkage applied with eBayes (robust = TRUE)^33^. Statistical significance was assessed using the Benjamini–Hochberg (BH) false discovery rate (FDR); no genes reached the pre-specified threshold of FDR < 0.05 in either tissue, consistent with the limited statistical power of n = 4 biological replicates after genome-wide multiple testing correction. For visualisation, volcano plots were generated with ggplot2 (v3.x) and ggrepel (v0.9.x)^34^, highlighting genes meeting a nominal threshold of p < 0.01 and |log₂ fold change| > 1 (CCR7^ΔcDC^ relative to CCR7^wt^).

### Bulk RNAseq data processing of sorted mouse co-cultured cDC

Wildtype cDC type 1 (cDC1; Live CD45⁺Ly6C⁻F4/80⁻CD19⁻CD3⁻CD11c⁺MHCII⁺XCR1^+^SIRPa^-^) and type 2 (cDC2; Live CD45⁺Ly6C⁻F4/80⁻CD19⁻CD3⁻CD11c⁺MHCII⁺XCR1^-^SIRPa^+^) were sorted after 18h of culture in complete RPMI media, 50 ng/ml Lipopolysaccharide (LPS), 10 ng/ml recombinant murine IL-10 (Biolegend) and 10 ng/ml recombinant murine TGF-beta 1 (Biolegend), co-culture with proliferating primary fibroblasts or senescent primary fibroblasts, or the combination treatments IL-10 and TGF-beta 1 or senescent fibroblasts 6 hours before adding LPS for 18h (n = 3). Total RNA was extracted as explained above. Strand-unspecific RNA-seq libraries were prepared and sequenced at the Genentech Sequencing facility as explained above. Raw sequencing reads were processed through the Roche internal Sunrise platform using the congee_cromwell_prd-ngs_pipeline workflow. Briefly, reads were quality-trimmed and adapter-clipped, then aligned to the mouse reference genome (GRCm38 / mm10, Ensembl release 90, Genomitory annotation GMTY230) using STAR. Gene-level read counts were quantified over exonic regions using HTSeqGenie (gene_exonic count type). Raw exonic read counts were imported into R (v4.5). Genes were retained if they passed expression filters using filterByExpr (edgeR v4)^30^ with default parameters applied per cell type to remove lowly expressed genes. Normalisation factors were computed with the trimmed mean of M-values (TMM) method^30^. For each gene, mean log₂-CPM values were computed across three biological replicates per condition (untreated, FB, senFB, LPS) in cDC1 and cDC2 separately.

PCA was performed on the 500 most variable genes (ranked by across-sample variance) from the log2-CPM matrix using prcomp with scaling (scale. = TRUE). Individual replicates are shown as coloured points labelled with replicate number; condition centroids are shown as open circles. Axes are displayed on equal scales (coord_fixed) to preserve geometric relationships between principal components. Percentage variance explained is indicated on each axis.

Differential expression between pairs of conditions was performed using limma-voom with a replicate-blocked design (∼ replicate + condition). Briefly, raw counts were passed to voom for mean-variance modelling, followed by lmFit and eBayes. Genes were considered differentially expressed at FDR < 0.05 and |log2FC| > 0.5 relative to the reference condition.

Unbiased, data-derived activatory and tolerogenic signatures were defined per cell type. For each cell type, differential expression was performed using limma-voom with a replicate-blocked design comparing (i) LPS versus untreated and (ii) tolerogenic versus untreated, using the DGEList restricted to the seven-condition set. Genes significantly upregulated relative to untreated (FDR < 0.05, log2FC > 0.5) were called significant genes per cell type, creating two gene sets per contrast . The intersection from both cell types was ranked by mean moderated t-statistic across cDC1+cDC2 and took the top 30 genes for visualization. Heatmaps for the four resulting signatures (activatory and tolerogenic, cDC1 and cDC2) were generated as described above, each restricted to its own cell type, using the seven-condition set, with genes ordered by descending z-score at the corresponding reference condition (LPS for activatory, tolerogenic condition for tolerogenic). Per-sample unbiased activatory and tolerogenic scores were computed as described above (Immunogenic and tolerogenic signature scores), using each cell type’s own unbiased gene lists, and displayed as an equal-scale scatter plot (“dotplot”) of unbiased tolerogenic score against unbiased activatory score, per cell type. All analyses were performed in R v4.5.

### Human scRNAseq public datasets analysis

Single-cell RNA sequencing data from the Human Lung Cell Atlas (HLCA) core dataset^23^ were obtained from the CellxGene repository (accession: 4cb45d80-499a-48ae-a056-c71ac3552c94), comprising 584,944 cells from 107 donors across 14 datasets. Pre-computed harmonized cell type annotations and UMAP embeddings were used as provided by the original authors without modification. Immune cells were identified using the top-level annotation (ann_level_1 == “Immune”). Conventional dendritic cell (cDC) subtypes — cDC1, cDC2, and Migratory DCs — were identified using the finest-resolution harmonized annotation (ann_finest_level). For each donor, the proportion of each cDC subtype relative to cDC. To characterize the maturation state of Migratory DCs, normalized gene expression values for canonical cDC maturation markers (CCR7, LAMP3, FSCN1, CD83, MARCKS, ITGAX, CLEC9A, CD1C) were visualized as a dot plot across cDC subtypes, displaying mean expression and percentage of expressing cells in stratified donors into two age groups: young (< 30 years) and aged (≥ 65 years).

All analyses were performed in R (v4.5.0) using the SingleCellExperiment, SummarizedExperiment, zellkonverter, dsassembly, Matrix, dplyr, and ggplot2^34^ packages.

### Luminex analysis

Serum cytokines were quantified using the Millipore Mouse Panel III Th17 25-plex panel. Briefly, serum samples were thawed, centrifuged 10 min at 10,000 g, diluted 1:10 in assay buffer, and analyzed following manufacturer’s instructions. Fecal immunoglobulins were quantified using Millipore Mouse Panel V IgG Isotyping 6-plex. Briefly, fecal samples were weighed, mechanically disaggregated in PBS containing protease inhibitors (Roche), centrifuged 10 min at 10,000 g twice, diluted in assay buffer, and analyzed following manufacturer’s instructions. Samples were acquired on a Luminex 200 (Thermo Fisher Scientific).

### Tissue processing and routine histology

Tissue samples were formalin-fixed, paraffin-embedded, and sectioned (approximately 4–5 µm thick sections). Alternatively, tissues were fixed in paraformaldehyde 4% at 4°C, OCT embedded and frozen, and sectioned (10–15 µm). For routine histological analysis, sections were stained with hematoxylin and eosin. Slides were scanned on the Nanozoomer S360 platform (Hamamatsu) at ×20 magnification.

Mouse tissues were first qualitatively assessed for morphologic changes and presence of immune aggregates, then they were randomly and blindly scored for inflammation according to the following matrix: 0: No notable inflammation; 1: Minimal inflammation, loose immune cells with no tissue disruption; 2: Mild inflammation; inflammatory aggregates with limited tissue architecture disruption; 3: Moderate inflammation; inflammatory aggregates with pronounced disruption of the tissue architecture; 4: Marked inflammation; coalescing to extensive inflammation disrupting the tissue. For colon tissue, mature aggregates were defined as aggregates with evident germinal centers.

Human samples were qualitatively scored for presence of lymphocyte/immune infiltrates as follows: 0: No infiltrates observed; 1: Minimal, ill-defined leukocyte infiltrates; 2: Small aggregates with minimal to no disruption of tissue architecture; 3: Mild to moderate aggregates with mild expansion.

### Immunohistochemistry and immunofluorescence

Immunohistochemistry was performed for CD3 (1:200 SP7 rabbit monoclonal antibody, Thermo Scientific), B220 (0.07 µg/ml RA3-6B2 rat monoclonal antibody, Pharmingen), IgG (0.25 µg/ml donkey anti-mouse IgG antibody with biotin, Jackson ImmunoResearch), and CLEC9A (1.9 µg/ml EPR22324 rabbit monoclonal antibody, Abcam). Antibodies were incubated for 60 minutes at 37 C, except for the IgG antibody, which was incubated at room temperature. CC1 standard retrieval (Roche) and bovine serum albumin as a blocking agent were used for CD3, B220, and CLEC9A. The CLEC9A assay also incorporated Ventana Inhibitor CM as an additional block. IgG antigen retrieval was performed with Target pH 6 solution, and non-specific labeling was blocked with hydrogen peroxidase and Sytek Biotin Block. CD3 was detected with the OmniMap detection system with diaminobenzidine (DAB) as the chromogen. B220 detection utilized a rabbit anti-rat linker antibody followed by Rabbit OmniMap detection with DAB. CLEC9A was detected with anti-HQ HRP (Roche) with DAB. The ABC-HRP Elite detection system with DAB (Vector Labs) was used for the IgG assay. Isotype matched antibodies were used as negative controls for each assay, and the specificities of the antibodies were confirmed in normal mouse or human tissues, based on the target species of the assay. Images were acquired using the Olympus VS200 Research Slide Scanner (Olympus Life Science). Whole slide scans were viewed using both the native viewer (OlyVIA) and the in-house Genentech gSlideViewer.

Immunofluorescence was performed on formalin-fixed, paraffin-embedded sections following antigen retrieval with Tris pH6 or EDTA pH9 (for intracellular antigens) or, alternatively, on OCT embedded frozen tissue sections. Sections were permeabilized in PBS containing 0.5% Triton X-100. Sections were blocked with Fish Serum Blocking Buffer (Thermo Fisher), then incubated with primary antibodies diluted in blocking buffer overnight at 4 °C. After washing with PBS-T, sections were incubated with fluorophore-conjugated secondary antibodies and/or streptavidin for 1 h at room temperature. Nuclei were counterstained with DAPI or live/death nuclear die, and slides were mounted with ProLong Gold Antifade mounting medium (Invitrogen). Images were acquired using a fluorescence confocal STED SP8 or Stelaris Nikon microscope and processed using identical settings across experimental groups. All antibodies are listed in Extended Data Table 6.

### RNA In Situ Hybridization (ISH)

Formalin-fixed, paraffin-embedded tissue sections (4-5 μm) were processed using the RNAscope Multiplex Fluorescent Reagent Kit v2 (Advanced Cell Diagnostics). Following hybridization, amplifications and secondary hybridization steps were performed for each probe: mouse probes included Mm-Zbtb46 (518313), Mm-Ccl19 (432881) and MmCxcl13 (407311). After the final amplification, the slides were washed and counterstained with DAPI, ProLong Gold mounted with Antifade mounting medium (Invitrogen) and imaged. Images were acquired using a fluorescence confocal Stelaris Nikon microscope and processed using identical settings across experimental groups.

### Image analysis and quantification

Human samples were qualitatively scored for CLEC9A immunolabeling as follows: 1: Few, scattered immunolabeled cells; 2: Minimally increased, individual immunolabeled cells; 3: Mildly increased cells in portal tracts and/or the parenchyma; and 4: Moderately increased labeling in portal tracts.

In human tissues, mature cDCs were identified by immunofluorescence as CCR7⁺ or LAMP3⁺ cells that were negative for CD3 and CD20, with assignment further supported by dendritic morphology. CCR7 and LAMP3 were selected as markers of the mature migratory cDC state because canonical cDC markers, including CD11c, were observed to be reduced during maturation in immunofluorescence staining.

Mouse samples were qualitatively scored for CD3 immunolabeling as follows: 1: Individual CD3 positive cells; 2: Aggregates of CD3+ cells that do not disrupt the tissue architecture; 3: Expansive aggregates of CD3+ aggregates that disrupt the tissue architecture; 4: Extensive CD3+ cellular infiltrates.

Image analysis of immunofluorescence stainings was performed using Fiji/ImageJ and QuPath.

### Laser Capture Microdissection (LCM)

Polypropylene microwell chips containing wells with a diameter of 2.2 mm were manufactured on polypropylene substrates by Protolabs1 (Maple Plain, MN, USA) and used for both sample capture and downstream proteomic processing. LC–MS-grade water, isopropanol, formic acid (FA), chloroacetamide (CAA), tris(2-carboxyethyl)phosphine hydrochloride (TCEP-HCl), and acetonitrile were obtained from Thermo Fisher Scientific (Waltham, MA, USA). N-dodecyl β-D-maltoside (DDM), HPLC-grade dimethyl sulfoxide (DMSO), sodium chloride (NaCl), and ammonium bicarbonate (ABC) were purchased from Sigma-Aldrich (St. Louis, MO, USA). Sequencing-grade trypsin was sourced from Promega (Madison, WI, USA).

For lymphoid aggregate proteome, 4 different lymphoid aggregates 2 different CCR7^ΔcDC^ mice were annotated in 5 serial cuts. For control tissue proteome, 4 different tissue areas from 2 different CCR7^ΔcDC^ mice were annotated in 5 serial cuts. Tissue regions of interest were isolated and collected using a PALM MicroBeam laser microdissection platform (Carl Zeiss MicroImaging, Munich, Germany). The instrument was equipped with a RoboStage to enable precise laser-guided micromanipulation and a RoboMover to facilitate transfer of dissected material. This setup allowed tissue fragments to be deposited directly into individual microwells of the microPOTS chips^35,36^. Before microdissection, each microwell was prepared with 1.6 μL of aqueous 50% DMSO buffer supplemented with 100 mM NaCl and 100 mM ABC. Dissected tissue material was collected into assigned microwells, with individual wells treated as technical replicates.

Following tissue collection, the microwell chip was heated at 95 °C for 2 h to promote decrosslinking. The chip was then incubated at 60 °C to remove DMSO by evaporation. Next, 1.5 μL of lysis and digestion solution was added to each well using an HP D100 nanoliter liquid dispenser. This solution consisted of 2 mM TCEP, 5 mM CAA, 0.05% DDM, and 5 ng/μL trypsin prepared in a 100 mM ABC buffer. Protein digestion was performed by incubating the chip at 37 °C for 10 h. After digestion, the resulting peptides were transferred onto Evotip Pure tips and desalted according to the manufacturer’s recommended protocol.

### LC–MS analysis

Peptide separation was performed on an Evosep system using the Whisper Zoom 40 samples-per-day method. Samples were separated on an IonOpticks Aurora Elite analytical column measuring 150 mm × 75 μm and packed with 1.7 μm particles. The analytical column was maintained at 60 °C during chromatographic separation.

Peptide analysis was conducted on an Orbitrap Astral mass spectrometer coupled with a FAIMS Pro interface (Thermo Fisher Scientific). The electrospray voltage was set to 1800 V, and the ion transfer tube temperature was maintained at 300 °C. FAIMS was operated at a compensation voltage of −50 V using user-defined mode, with the outer electrode temperature set to 90 °C. Data were acquired using a data-independent acquisition strategy. MS1 precursor scans were collected in the Orbitrap over a mass range of 400–900 m/z at a resolution of 240,000, with a normalized AGC target of 500% and a maximum injection time of 246 ms. MS2 fragment spectra were acquired in the Astral analyzer over a scan range of 150–2000 m/z, with the default precursor charge state set to 2. Precursor ions were fragmented using HCD at a normalized collision energy of 25%. The DIA isolation window was set to 5 m/z, with a maximum injection time of 10 ms and a normalized AGC target of 800%. Loop control was time-dependent, with a cycle time of 0.6s.

### Proteomic data analysis

Database: The Swiss-Prot protein sequence database of Mus musculus (downloaded in 2024 May, containing 22092 reviewed protein sequences) was used for database searching.

Raw mass spectrometry data were analyzed in Spectronaut version 20.2 using the directDIA workflow. Default processing parameters were used unless otherwise noted, with cross-run normalization turned off. The database search was configured for Trypsin/P specificity and permitted up to two missed cleavages. Peptides ranging from 7 to 52 amino acids in length were considered. Oxidation of methionine and acetylation at the protein N-terminus were included as variable modifications, whereas cysteine carbamidomethylation was specified as a fixed modification. Identification results were controlled at a false discovery rate of <1% at the peptide-spectrum match, peptide, and protein levels. To maximize identifications, the deep directDIA+ search option was used within the directDIA analysis workflow. Protein quantification was restricted to proteotypic peptides by applying the “Only Proteotypic” setting for proteotypicity filtering, thereby excluding peptide sequences shared across multiple proteins. The experiment-level q-value cutoff was set to 0.01 for both precursors and proteins, and the precursor posterior error probability cutoff was set to 0.2. At the run level, the protein q-value cutoff was set to 0.05, while the protein posterior error probability cutoff was set to 0.75. Quantification was performed using MS1-level signal intensity.

The precursor-level report generated by Spectronaut was subsequently imported into MSstats version 4.12.0 for downstream processing, including equalized median normalization, missing precursor imputation, and protein summarization.

### Spatial Proteomics Data Processing and Differential Protein Analysis

Protein intensity data from spatial proteomics profiling of lymphoid structures isolated from lung, salivary gland, and liver, together with matched control tissues (stromal tissue from wild-type tissue from the same organs), were processed in Python (v3.12) using pandas (v2.x) and NumPy. Raw intensity values were treated as label-free quantification (LFQ) intensities, with missing values (NaN) interpreted as below-detection-limit (not detected).

For each organ independently, proteins were classified as exclusively detected in lymphoid structures if they were quantified (non-missing) in at least two of four lymphoid structures biological replicates and were completely absent (all values missing) across all control replicates of the same organ (wild-type tissue, n = 4).

For each organ, proteins detected in at least two replicates in both the TLS and control groups were tested for differential abundance using a two-sided Welch’s t-test applied to log₂-transformed intensities (log₂[intensity + 1]). Multiple testing correction was performed using the Benjamini–Hochberg false discovery rate (FDR) procedure. Proteins with an absolute log₂ fold-change > 1 (i.e., > 2-fold higher in TLS) and an FDR-adjusted p-value < 0.05 were considered significantly upregulated in lymphoid structures.

For each organ, the lymphoid structures-enriched protein set was defined as the union of exclusively detected and significantly upregulated proteins. Four gene set collections were defined for downstream enrichment analysis: (i) per-organ lymphoid structures sets (liver, lung, salivary gland); (ii) a core lymphoid structures signature comprising proteins present in the lymphoid structures -enriched sets of all three organs (intersection); and (iii) a pan-lymphoid structures union comprising proteins enriched in lymphoid structures in at least one organ.

### Pathway and Gene Ontology Enrichment Analysis

Over-representation analysis (ORA) was performed for each gene set using the Enrichr API accessed via the gseapy Python package (v1.3.0). Four curated gene set databases were queried independently: Gene Ontology Biological Process (GO BP 2023), KEGG Mouse 2019, Reactome 2022, and MSigDB Hallmark 2020. Mouse gene symbols were submitted directly to Enrichr using the mouse organism setting. Enrichment significance was assessed using Fisher’s exact test implemented within Enrichr, and results were corrected for multiple testing using the Benjamini–Hochberg FDR method. Pathways with FDR-adjusted p-value < 0.05 were considered statistically significant. The overlap ratio was defined as the number of query genes annotated to a given pathway divided by the total number of genes in that pathway.

### Visualization

Results were visualized in Python using matplotlib. For each database and query group, dot plots were generated displaying the top 20 enriched pathways ranked by -log₁₀(FDR), with dot size proportional to the overlap ratio and dot colour encoding -log₁₀(FDR) intensity (YlOrRd colormap). Pathway enrichment across all query groups was additionally summarised as heatmaps (-log₁₀FDR matrix) and bubble plots. Protein overlap across organs was visualised using three-way Venn diagrams (matplotlib-venn). Volcano plots were generated for each organ displaying log₂ fold-change (TLS vs. control) against -log₁₀(FDR), with significantly upregulated proteins (log₂FC > 1, FDR < 0.05) highlighted.

Key software versions: Python 3.12; pandas; NumPy; SciPy; statsmodels (Benjamini–Hochberg correction); gseapy 1.3.0; matplotlib; matplotlib-venn.

### Quantification, statistical analysis and data presentation

Data analysis was performed as noted in the figure legends. Statistical analysis and data presentation were performed using GraphPad Prism 10.1.0. P values are reported.

### Statistical analysis

GraphPad Prism 10 (GraphPad Software, v10.0.2) was used for most bar graph visualizations and to calculate significant differences. Unless otherwise indicated, each data point represents an independent biological replicate (individual mouse or human donor). Multiple tissue regions or lymphoid aggregates obtained from the same biological specimen are identified as nested measurements and were not treated as independent biological replicates. For in vitro studies, replicates indicate independent biological samples randomly allocated into experimental groups. For in vivo studies, replicates indicate individual mice allocated randomly to experimental groups. No statistical methods were used to predetermine sample size. Sample sizes were based on prior experience with the experimental models and were sufficient to detect reproducible biological differences across independent experiments. Exact numbers of biological replicates are reported for each experiment in the figure legends. All statistical comparisons are two sided. Difference testing between groups was done using t tests, one-way or two-way ANOVA followed by multiple comparisons as indicated in the figure legends for each experiment.

P values are indicated in the corresponding figures or, for body-weight change curves, in the associated raw data files.

## Data availability

The Human Lung Cell Atlas (HLCA) core dataset analysed in this study is publicly available from the CellxGene data portal (collection ID: 4cb45d80-499a-48ae-a056-c71ac3552c94; https://cellxgene.cziscience.com) and from Zenodo (https://doi.org/10.5281/zenodo.6337966), as described in the original study^23^. RNA sequencing data generated from this study will be available in the GEO repository upon publication. The mass spectrometry raw data will be available in the ProteomeXchange Consortium via the MassIVE partner repository upon publication. During peer review, the deposited datasets will be kept private and made accessible to editors and reviewers using repository-provided reviewer credentials.

All other data supporting the findings of this study are available within the Article, Extended Data and Source Data files. Any additional information is available from the corresponding author upon reasonable request.

## Code availability

Analysis scripts used to generate the computational results and figures will be made publicly available upon publication.

## AI/LLM disclosure

Generative AI tools were used to assist with code development and/or language editing. All analyses, outputs and manuscript content were independently reviewed and validated by the authors.

## Authors Contributions

I.M. conceived the study, designed and performed the experiments, analysed and interpreted the data, and wrote and revised the manuscript. B.M.M. contributed to experiments, data analysis and interpretation, and provided critical input. B.O.J. contributed to computational analyses and interpretation, and provided critical input. J.P. performed immunohistochemistry experiments. K.M. contributed to the analysis of DSS colitis data. Z.Y. performed proteomic analyses, with supervision from Y.Z. J.W. analysed and interpreted histopathological data and provided expert pathology input. C.M. conceived and supervised the study, contributed to experimental design and data interpretation, and wrote and revised the manuscript. All authors reviewed and approved the manuscript.

## Acknowledgements

We thank the members of the Moussion lab and the Immunology department for their advice, discussions, and reagents. We thank multiple contributors from the Research Pathology Department Core Labs at Genentech, including Rae Liu, Linda Rangell, Margaret Solon, Wennie Chen, Deb Dunlap, Serena Lee, Treshan Myers, Mario Cabrera, and Jian Jiang for their support with histology and immunochemistry methods, Will Eart and Jeanette Villareal-Kinney for their help with tissue procurement, as well as the Digital Pathology Imaging group for their technical support. We acknowledge the Genentech mouse husbandry team, including Shannon Powell, Tina Scholl, Ben Torres, and Michael Long. We thank the veterinary care team and IACUC office members, including Emmanuel Chua, Millicent Lu, Gaby Fuentes, and Catherine Sohn, for their dedicated animal care and support. We also appreciate the support of the Genentech Laboratory Animal Resources team, including Bridget Maguire-Colton, Caroline Sepulveda, Dorie Montoya, Ryan Scott, and Bruno Alicke. We acknowledge the Genentech Flow Cytometry Core group for technical assistance, including Terence Ho, George Tweet, CK Poon, Alice Tan, Cole Meyers, Kevin Le, Jonathan Paw, and Jovencio Borneo. We are grateful to the Genentech Center for Advanced Light Microscopy (CALM) core group, including Melissa Gonzalez Edick, David DeWitt, and Sarah Gierke, for imaging support. We thank Yuxin Liang, Ana Magalhaes and Marco De Simone for assistance with RNA sequencing data pipelines. Finally, we acknowledge the Genentech Postdoctoral Program for its support, particularly Alexandra Colon-Rodriguez, Patsy Jaboneta, Molly Taylor, Vishva Dixit, and Saptarsi Haldar. This study was funded by Genentech, a member of Roche.

---

**Extended Data Fig. 1.**
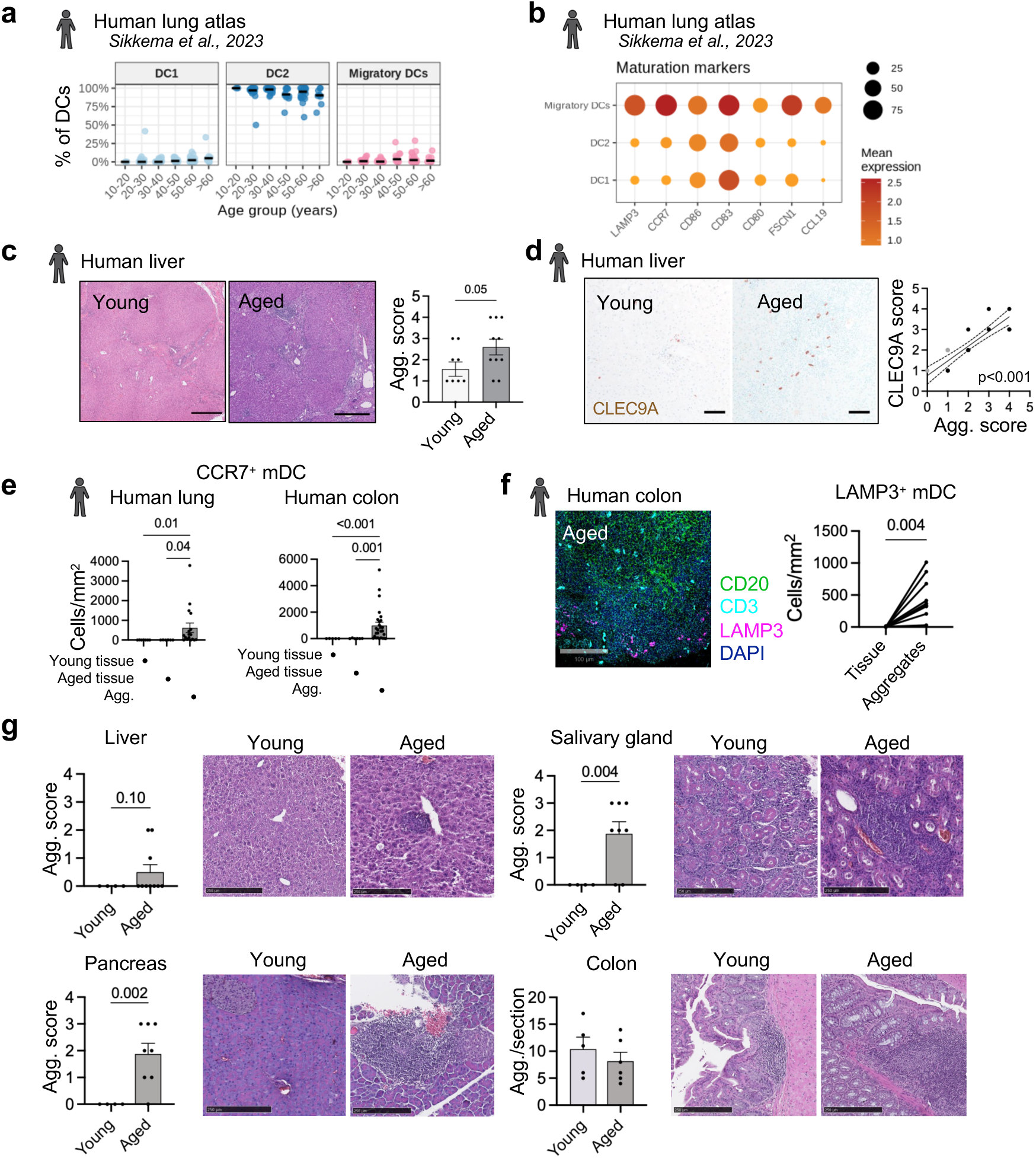
Mature cDCs associate with lymphoid aggregates across human and mouse aged tissues. a, Proportion of conventional dendritic cells (cDCs) across ages in a published human-lung scRNA-seq atlas dataset^23^. **b,** Expression of markers defining mature cDCs in the same dataset. **c,d,** Representative H&E images and lymphoid-aggregate scores (**c**) and CLEC9A immunohistochemistry with correlation between CLEC9A+ cDC and aggregate scores (**d**) in young (<50 years; gray points; *n* = 9) and aged (>70 years, black points; *n* = 10) human liver presenting lymphoid aggregates. Two-tailed unpaired *t*-test (**c**); linear regression and two-tailed Spearman correlation (r = 0.8916) (**d**). Scale bars 500 µm (H&E) and 200 µm (IHC). **e,** CCR7+CD20-CD3-mature cDC abundance in tissue parenchyma of young (*n =* 7 lung, 5 colon) and aged (*n =* 6 lung, 6 colon) human tissues and lymphoid aggregates in aged tissues (*n* = 17 aggregates from 6 lung samples, 27 aggregates from 6 colon samples). One-way ANOVA with two-stage step-up correction. **f,** Representative immunofluorescence and quantification of LAMP3+CD20-CD3-mature cDCs in parenchyma and aggregates from young (*n* = 5) and aged colon (*n* = 5). Two-tailed paired t test. Scale bars 100 µm. **g,** Histological and immunofluorescence analysis of lymphoid aggregates in young (*n* = 4 liver, 4 salivary gland, 4 pancreas, 5 colon) versus aged mouse (*n* = 10 liver, 8 salivary gland, 8 pancreas, 6 colon) tissues under steady-state. Two-tailed unpaired *t*-test. Scale bars 250 µm. Data are mean ± s.e.m.

**Extended Data Fig. 2.**
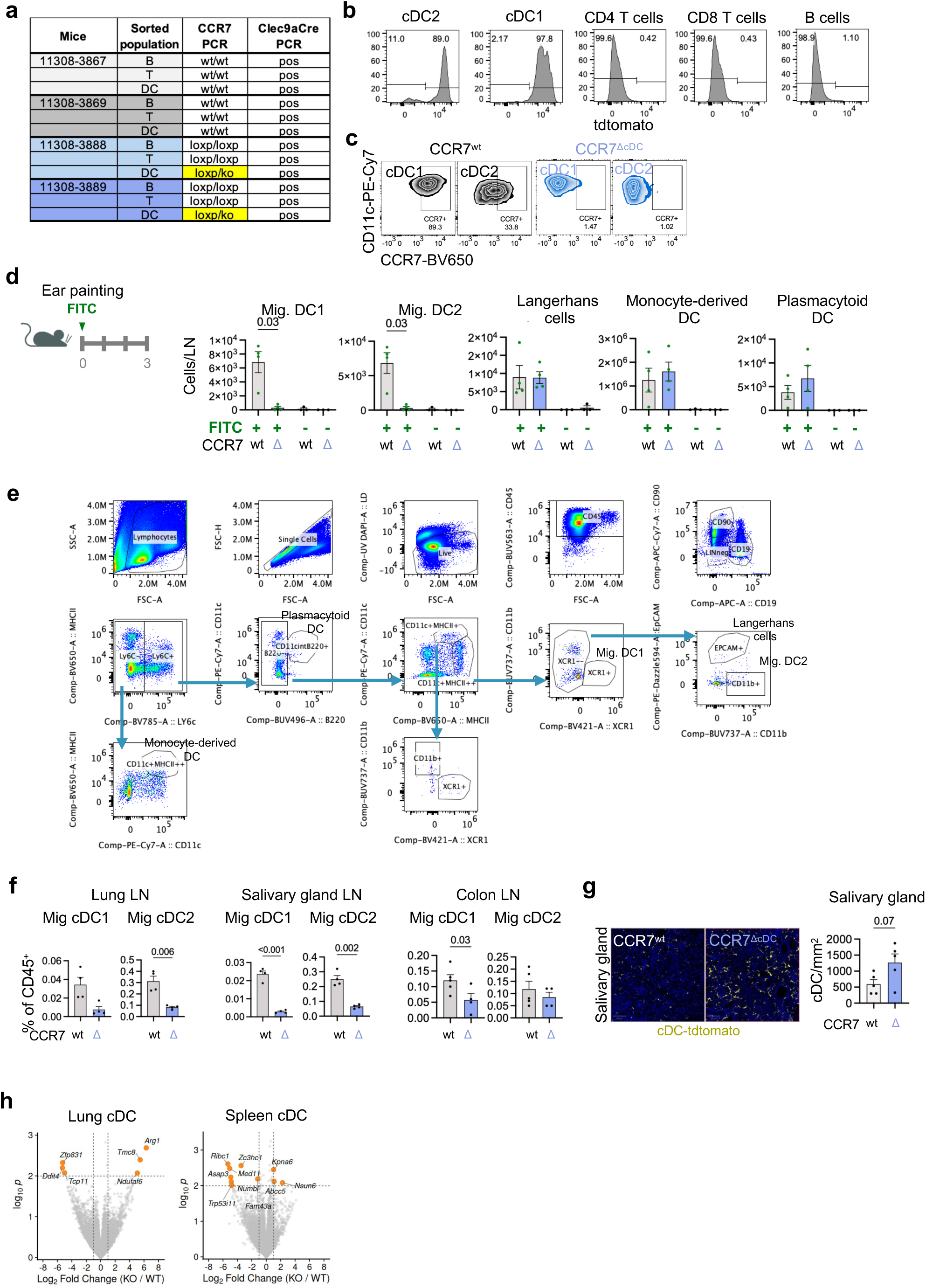
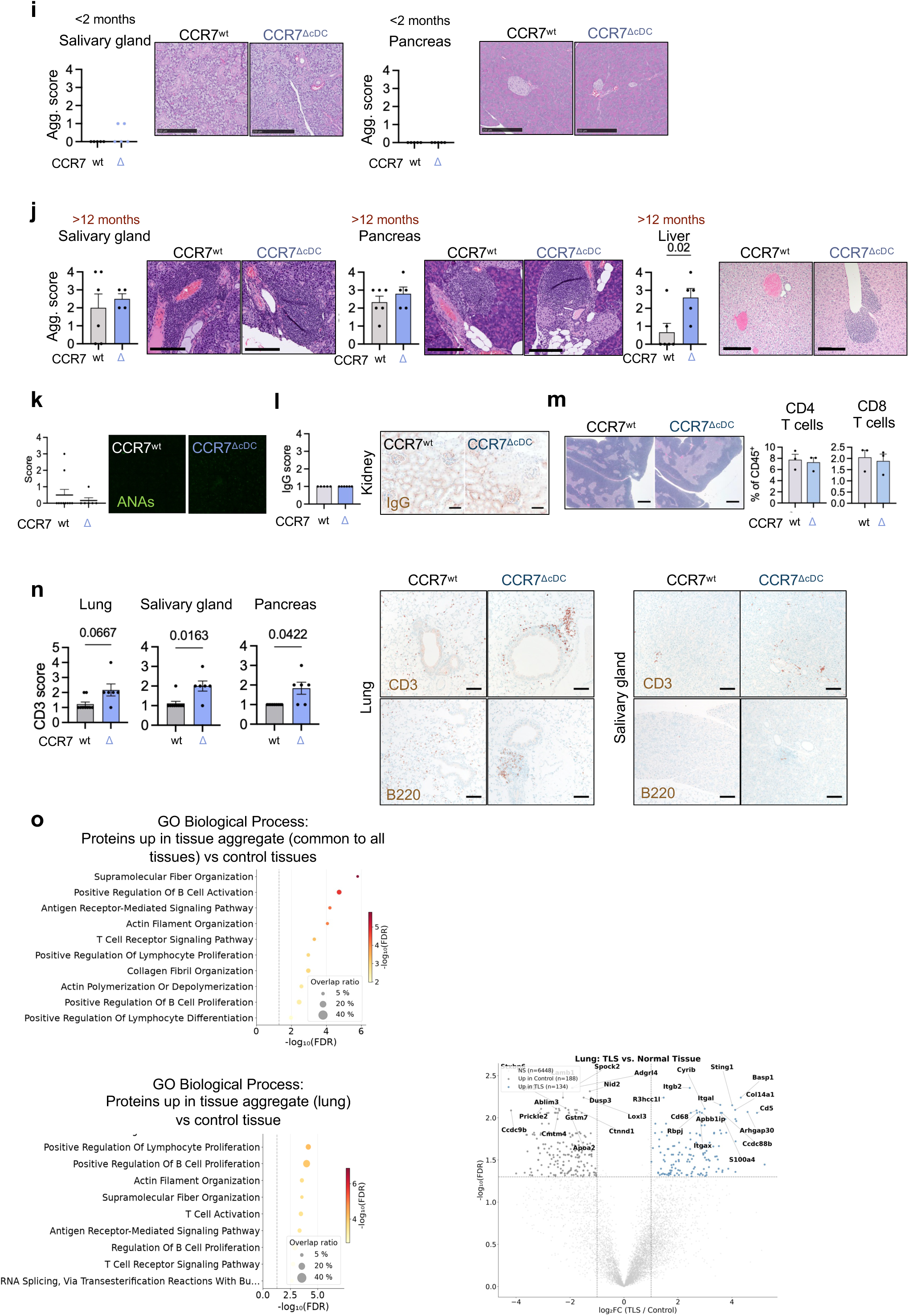
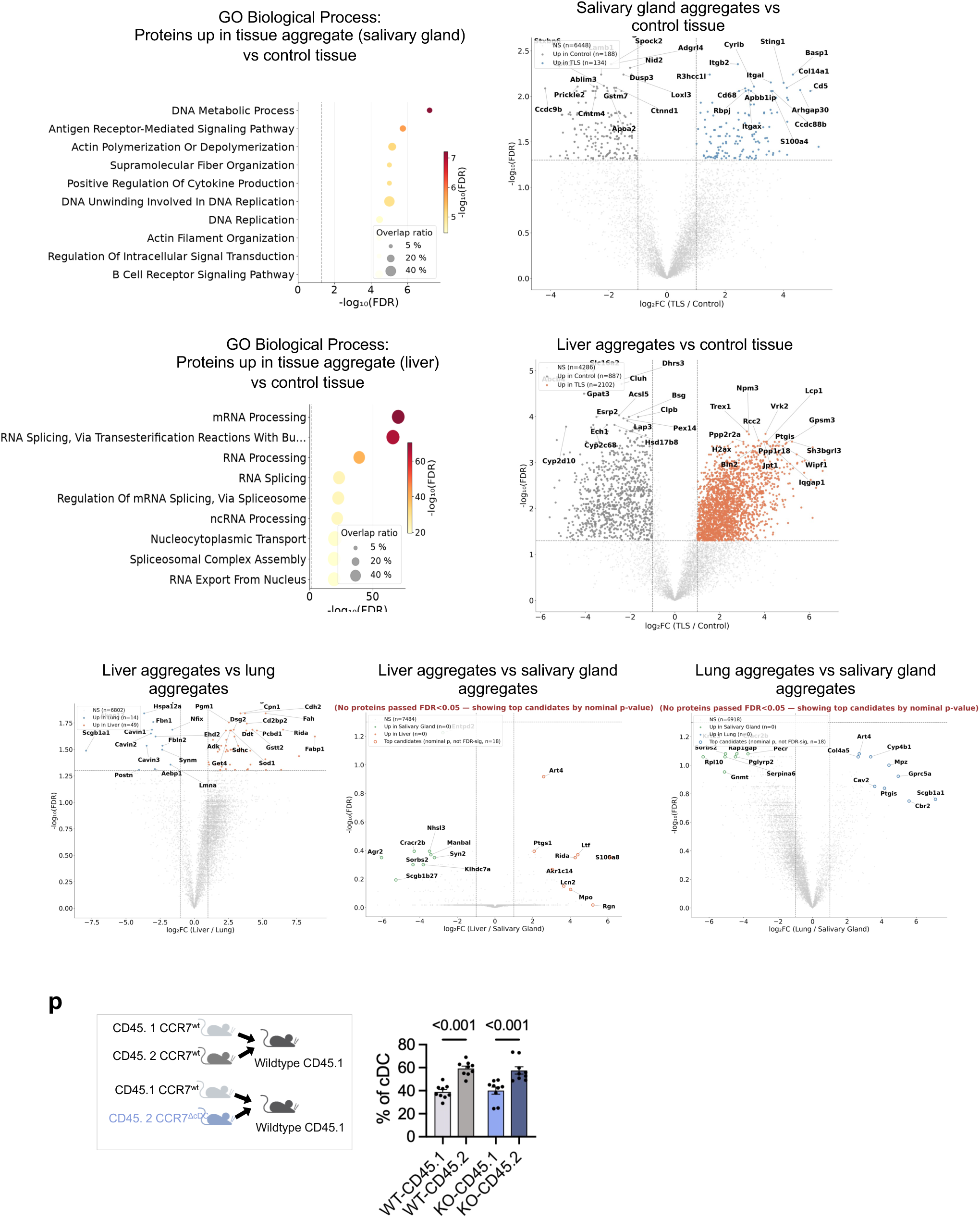
Validation and characterization of the CCR7^ΔcDC^ model. a, PCR analysis of CCR7-locus recombination and Cre expression in sorted splenic B cells, T cells and cDCs from CCR7^wt^ (*n* = 2) and CCR7^ΔcDC^ (*n* = 2) mice. **b,** Flow-cytometric analysis of tdTomato lineage tracing in splenic cDC1, cDC2, CD4+ T cells, CD8+ T cells and B cells. **c,** Flow-cytometric analysis of CCR7 expression in splenic cDCs from CCR7^wt^ and CCR7^ΔcDC^ mice after LPS stimulation. **d,** FITC ear-painting assay and quantification of FITC+ dendritic cells in draining lymph nodes from painted ears (FITC+; *n* = 4) and control unpainted ears (FITC-; *n* = 3) from CCR7^wt^ and CCR7^ΔcDC^ mice. One-way ANOVA with two-stage step-up correction. **e,** Gating strategy for cDC analysis in mouse lymph nodes **f,** Migratory cDC type 1 and 2 among CD45+ cells in tissue-draining lymph nodes from CCR7^wt^ (*n* = 4 lung, 4, salivary gland, 5 colon) and CCR7^ΔcDC^ (*n* = 4 lung, 4, salivary gland, 4 colon) mice. Two-tailed unpaired *t*-test; representative of two independent experiments. **g,** Representative immunofluorescence and quantification of tdTomato+ lineage-traced cDCs in salivary glands from *Clec9a*^Cre^;*Rosa26*^LSL-tdTomato^;CCR7^WT^ (*n* = 5) and CCR7^ΔcDC^ mice (*n* = 5). Representative of 2 independent experiments. Two-tailed unpaired *t*-test. Scale bars 50 µm. **h,** Differential gene-expression analysis of cDCs from CCR7^ΔcDC^ and littermate CCR7^wt^ mice. **i,j,** Representative H&E images and lymphoid-aggregate quantification of tissues from young (<2 months; *n* = 5 CCR7^wt^, 5 CCR7^ΔcDC^; **i**) and aged (>12 months; *n* = 6 CCR7^wt^, 5 CCR7^ΔcDC^, 4 CCR7^ΔcDC^ for salivary gland; **j**) CCR7^wt^ and CCR7^ΔcDC^ mice (*n* = 6). Scale bars 250 µm (h) and 200 µm (i).| **|k,** Serum antinuclear antibodies in CCR7^wt^ (*n* = 10) and CCR7^ΔcDC^ (*n* = 6) mice and quantification. **l,** and renal IgG deposition in CCR7^wt^ (*n* = 5) and CCR7^ΔcDC^ (*n* = 6) mice and quantification. Scale bars 50 µm **m**, Representative H&E images and proportion of CD4 and CD8 single positive T cells in the thymus of young <2 months CCR7^wt^ (*n*= 3) and CCR7^ΔcDC^ (*n*= 3) mice. Two-tailed unpaired *t*-test. Scale bars 500 µm. **n,** Quantification of CD3+ T-cell abundance and representative CD3 and B220 immunohistochemistry in CCR7^wt^ (*n* = 9) and CCR7^ΔcDC^ tissues (*n* = 6). Two-tailed unpaired *t*-test. Scale bars 100 µm **o,** Gene Ontology Biological Process enrichment of proteins selectively detected or increased in lymphoid aggregates from CCR7^ΔcDC^ tissues versus similar regions in CCR7^wt^ tissues and volcano plots representing genes up and down for the indicated comparisons. **p,** Bone-marrow chimera design and splenic contribution of CD45.1+ and CD45.2+ cDCs in recipient CD45.1 mice receiving bone marrow from CD45.1 CCR7^wt^ and CD45.2 CCR7^wt^ (*n* = 9) or CD45.1 CCR7^wt^ and CD45.2 CCR7^wt^ (*n* = 9) mice. One-way ANOVA with two-stage step-up correction. Data are mean ± s.e.m. where applicable.

**Extended Data Fig. 3.**
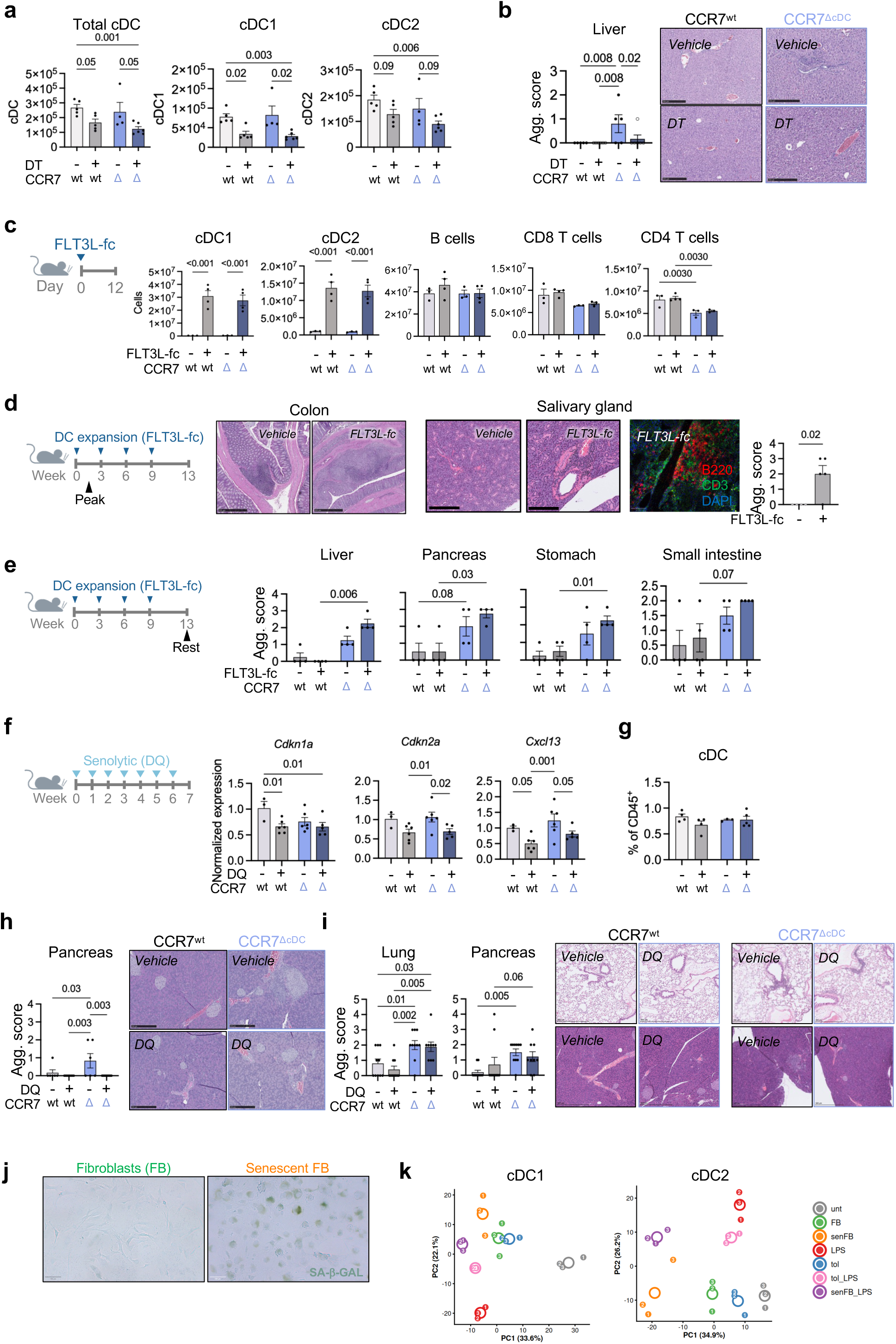
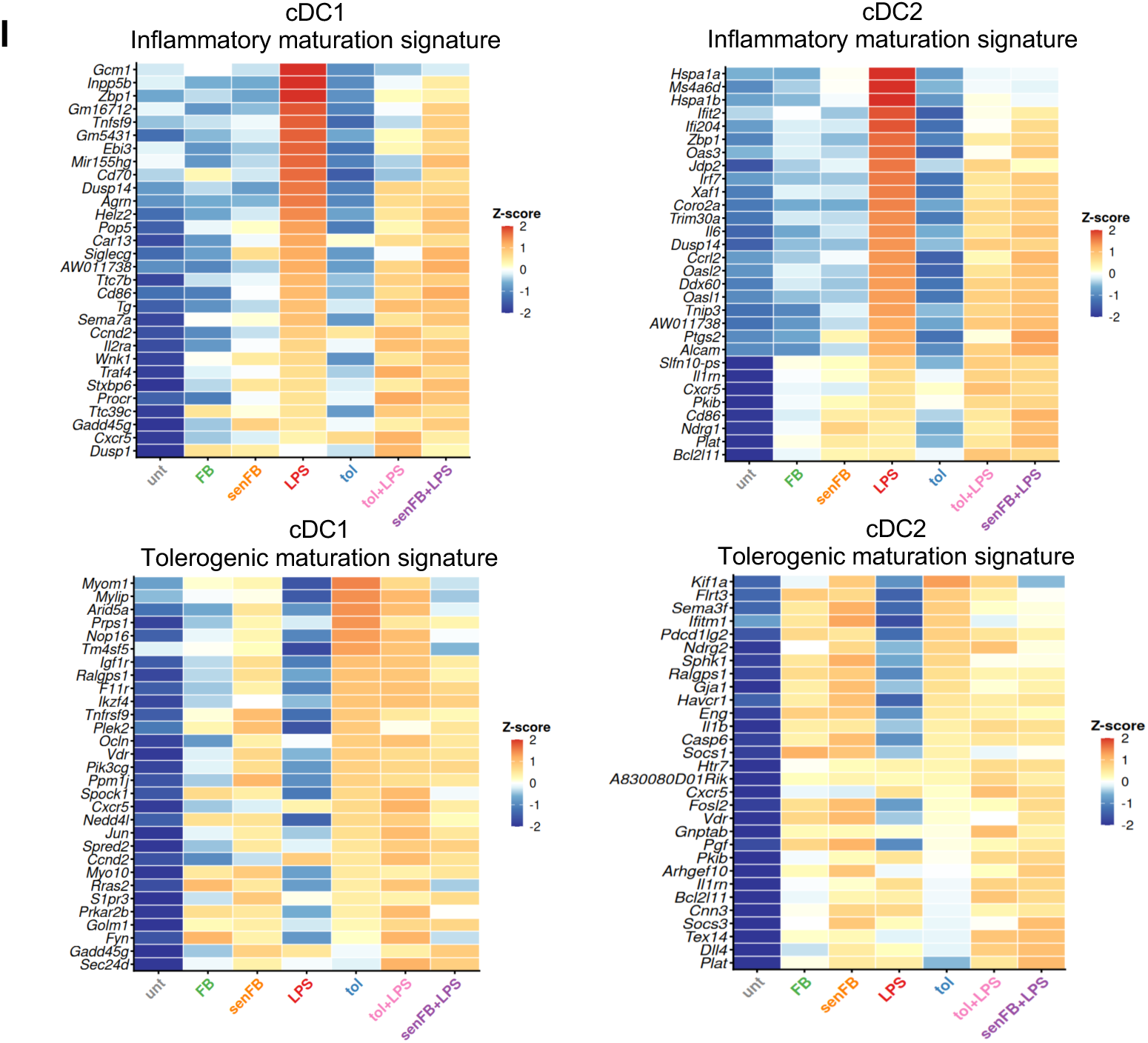
Experimental manipulation of tissue senescence, cDC abundance and cDC persistence. a, Splenic cDC abundance and lymphoid aggregates following 1 week of diphtheria toxin (DT)-mediated cDC depletion in CCR7^wt^;*Zbtb46*^DTR^ (*n* = 5 untreated, 6 treated) and CCR7^ΔcDC^;*Zbtb46*^DTR^ (*n* = 4 untreated, 6 treated) mice. One-way ANOVA with two-stage two-step correction. **b,** Representative H&E images and aggregate quantification in CCR7^wt^;*Zbtb46*^DTR^ (*n* = 5 untreated, 6 treated) and CCR7^ΔcDC^;*Zbtb46*^DTR^ (*n* = 5 untreated, 6 treated) mice untreated or after 1 week of DT-mediated cDC depletion. One-way ANOVA with two-stage two-step correction. Scale bars, 500 µm. **c,** Schematic of treatment and splenic immune-cell abundance and lymphoid aggregates 12 days after FLT3L–Fc treatment (peak of cDC expansion) (*n* = 3 untreated CCR7^wt^, 3 untreated CCR7^ΔcDC^, *n* = 4 treated CCR7^wt^, 3 treated CCR7^ΔcDC^). One-way ANOVA with two-stage two-step correction. **d,** Schematic of treatment and representative H&E and immunofluorescence images and aggregate quantification in wild-type adult mice following FLT3L–Fc-induced cDC expansion at peak expansion (day 12) (n = 4 untreated; n= 5 treated). Immunofluorescence shows B220+ B cells, CD3+ T cells and DAPI. Two-tailed *t*-test. Scale bars, 500 µm (colon H&E), 200 µm (salivary gland H&E) and 50 µm (immunofluorescence). **e,** Schematic of treatment and lymphoid-aggregate analysis after chronic FLT3L–Fc treatment, 30 days after the final treatment (rest) in CCR7^wt^ and CCR7^ΔcDC^ mice (*n* = 4 untreated CCR7^wt^, 4 treated CCR7^wt^, 4 untreated CCR7^ΔcDC^, 4 treated CCR7^ΔcDC^). One-way ANOVA with two-stage two-step correction. **f,** Schematic of treatment with vehicle or dasatinib plus quercetin (DQ). Lung expression of *Cdkn2a*, *Cdkn1a* and *Cxcl13* in CCR7^WT^ and CCR7^ΔcDC^ mice treated with vehicle or DQ for 7 weeks. One-way ANOVA with Fisher’s LSD test. **g,** Splenic cDC abundance in CCR7^WT^ and CCR7^ΔcDC^ mice treated with vehicle or DQ for 7 weeks. One-way ANOVA with two-stage two-step correction. **h,** Lymphoid-aggregate quantification in 2-month-old CCR7^WT^ and CCR7^ΔcDC^ mice treated with vehicle or DQ (*n* = 3–5). One-way ANOVA with two-stage step-up correction. Scale bars 250 µm. **i,** Lymphoid-aggregate quantification in 8-month-old (**e**) CCR7^WT^ and CCR7^ΔcDC^ mice treated with vehicle or DQ (*n* = 3–9). One-way ANOVA with two-stage two-step correction. Scale bars 500 µm. **j,** Senescence-associated beta galactosidase staining of control fibroblasts and senescent fibroblasts. Scale bars, 100 µm **k,** Principal-component (PC) analysis of the 500 most variable genes from TMM-normalized log2-CPM profiles in cDC1 (*n* = 3) and cDC2 (*n* = 3) across culture conditions. FB, fibroblast; senFB, senescent fibroblast; LPS, lipopolysaccharide; tol, tolerogenic factors. **l,** Heatmap of mean z-scored log2 counts per million (CPM) for cDC inflammatory and tolerogenic maturation signature genes in cDC1 (*n* = 3) and cDC2 (*n* = 3) across the same culture conditions. Data are mean ± s.e.m. where applicable.

**Extended Data Fig. 4.**
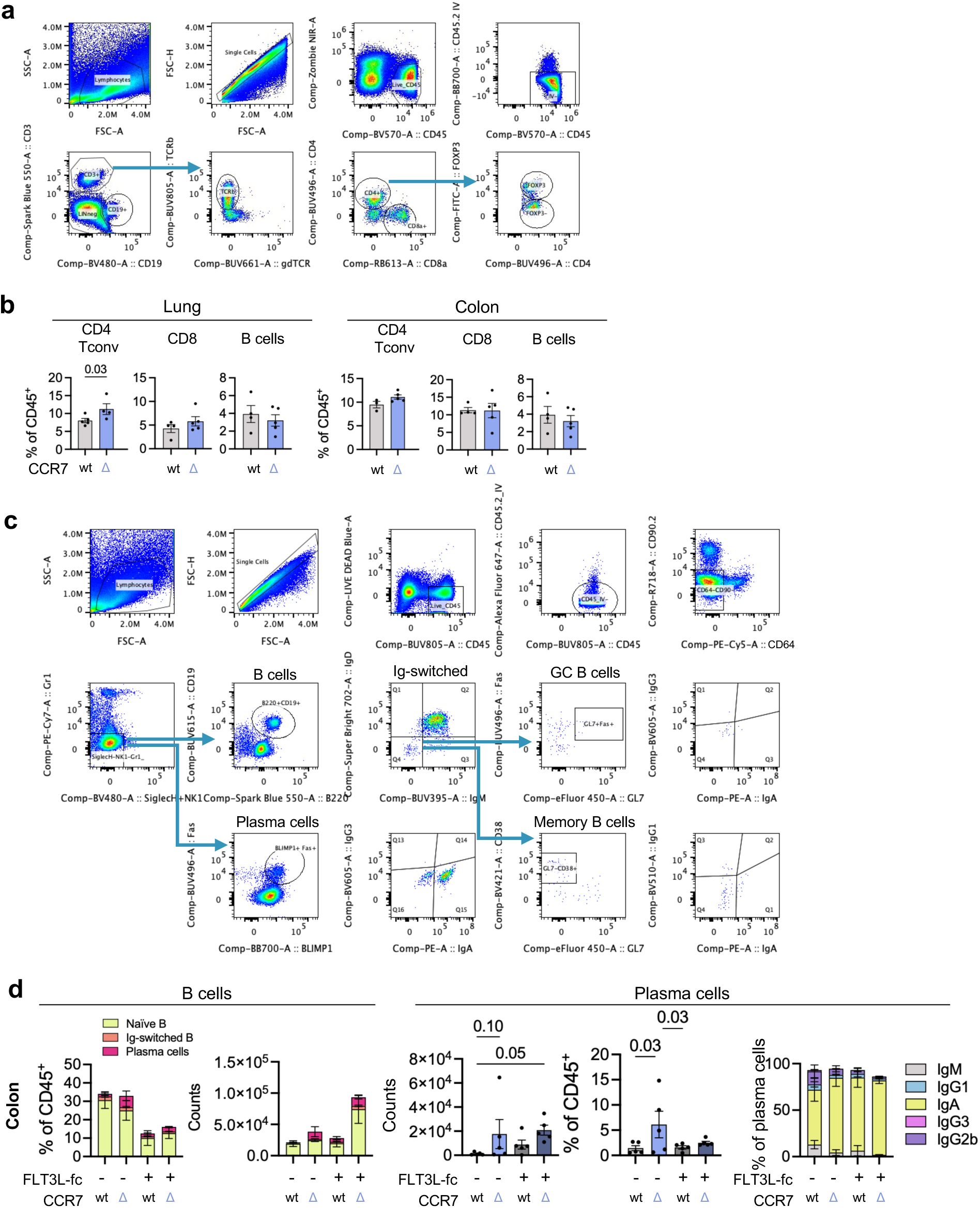
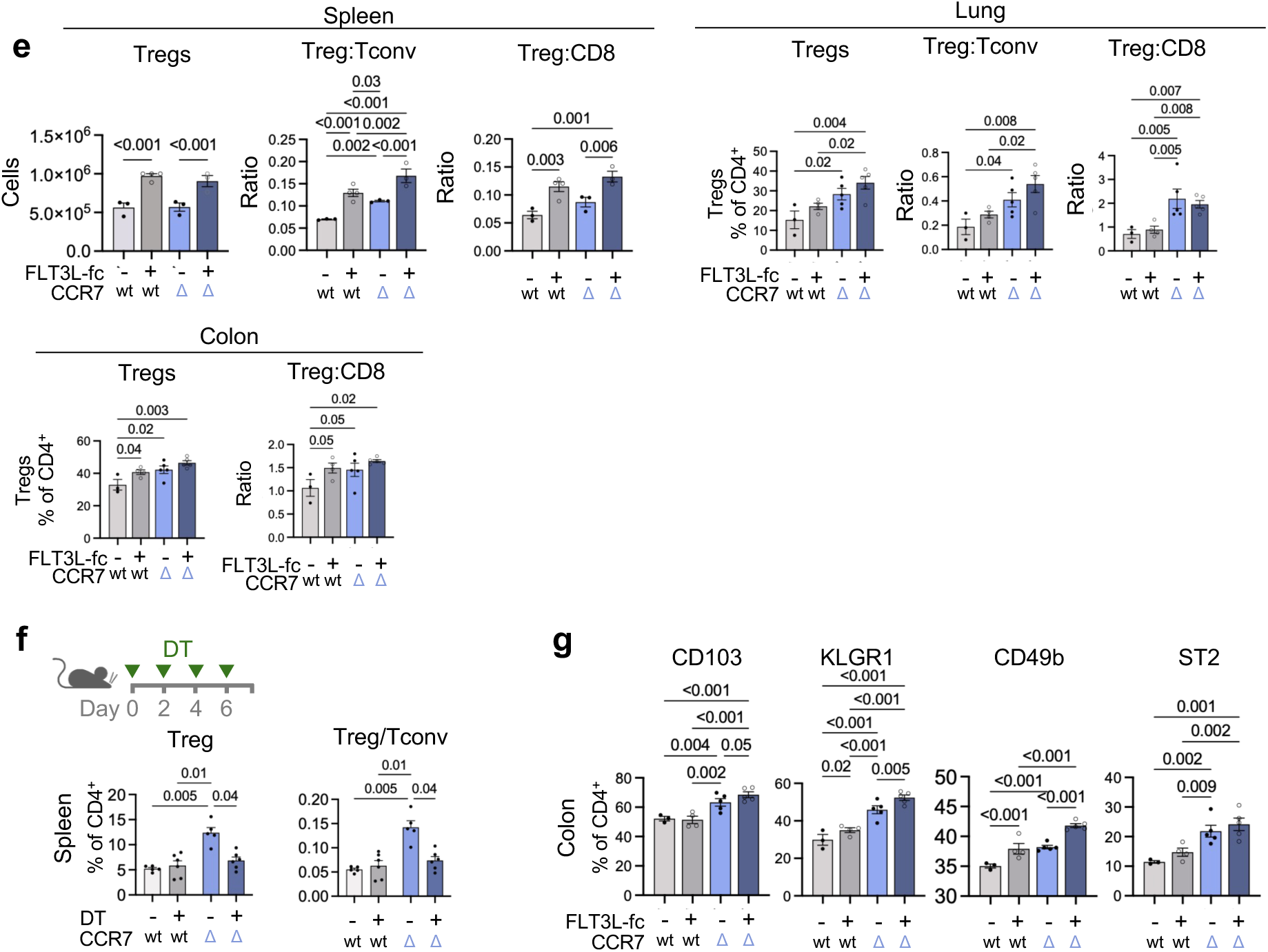
Lymphocyte composition following altered cDC persistence or abundance. a, Gating strategy for flow-cytometric analysis of T cells in mouse tissues. **b,** CD8+ T cells, CD4+ conventional T (Tconv) cells, Tregs and B cells as percentages of CD45+ cells in peripheral tissues from CCR7^WT^ and CCR7^ΔcDC^ mice (*n* = 3–5). Two-tailed unpaired *t*-test. **c,** Gating strategy for B-cell analysis. **d,** Flow-cytometric quantification of naive B cells, IgD−IgM− class-switched B cells and plasma cells and plasma-cell immunoglobulin isotypes in colon from untreated or FLT3L–Fc-treated CCR7^WT^ (*n* = 5 untreated and 5 treated) and CCR7^ΔcDC^ mice (*n* = 5 untreated and 5 treated). One-way ANOVA with Fischer’s LSD test. **e,** Treg burden as percentages of CD45+ and absolute numbers, Treg:CD8+ T-cell ratios and Treg:CD4 Tconv ratios in tissues from untreated or FLT3L–Fc-treated CCR7^WT^ and CCR7^ΔcDC^ mice tissues (*n* = 3–5). One-way ANOVA with two-stage step-up correction. **f,** Splenic Treg-cell abundance and Treg/Tconv ratio in CCR7^WT^;*Zbtb46*^DTR^ and CCR7^ΔcDC^;*Zbtb46*^DTR^ mice after 1 week of diphtheria toxin (DT)-mediated cDC depletion (*n* = 5–6). One-way ANOVA with two-stage step-up correction. **g,** Flow-cytometric expression of CD103, KLRG1, CD49b and ST2 by colon Treg cells from untreated and FLT3L–Fc-treated CCR7^WT^ and CCR7^ΔcDC^ mice (*n* = 3–5). One-way ANOVA with two-stage step-up correction. Data are mean ± s.e.m.

**Extended Data Fig. 5.**
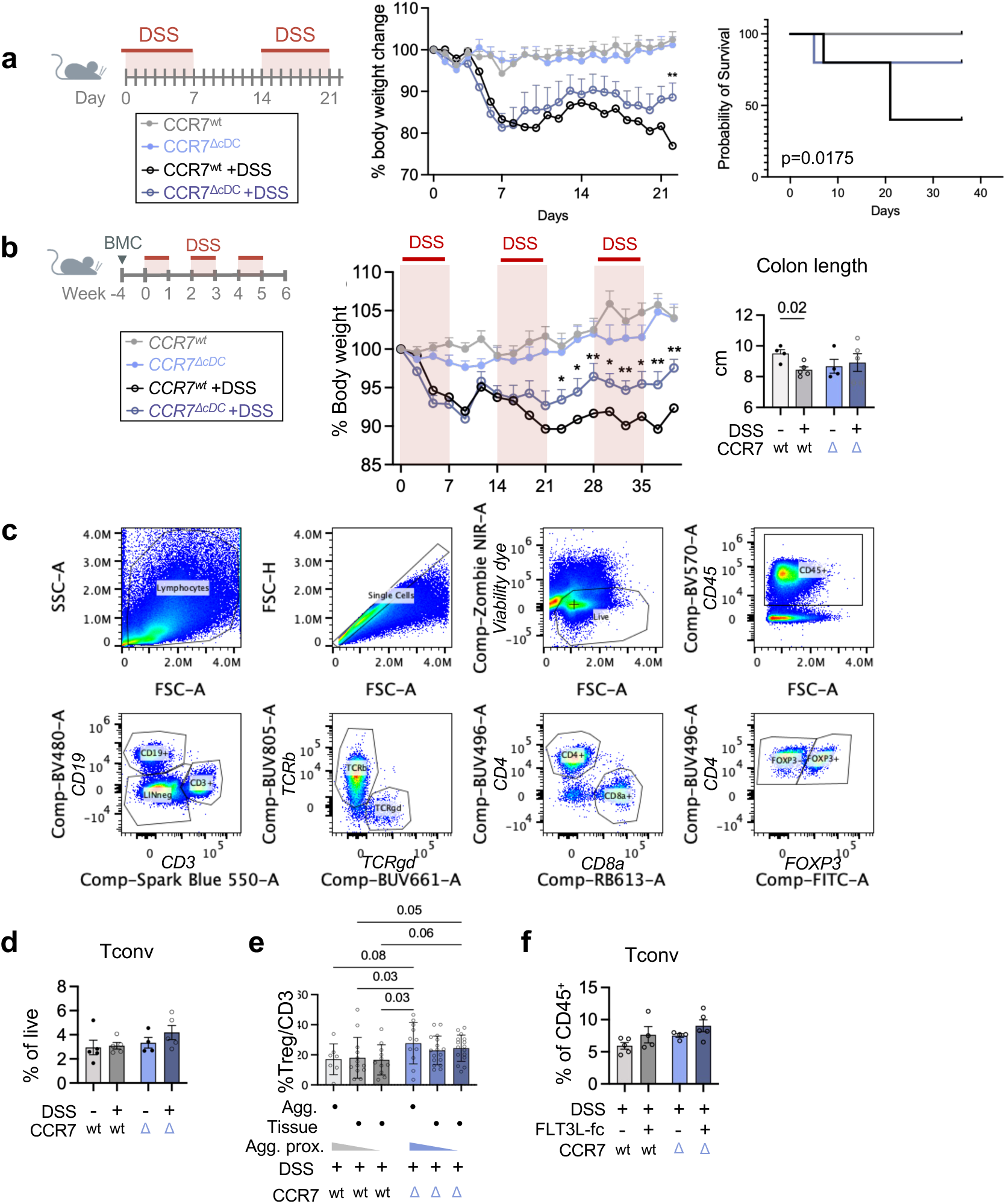
cDC persistence and expansion during experimental colitis. a, Chronic colitis induced by two 7-day cycles of 2.5% DSS separated by a 7-day recovery period, with longitudinal body weight and survival in CCR7^WT^ (*n* = 5 vehicle, 4 DSS) and CCR7^ΔcDC^ (*n* = 5 vehicle, 4 DSS) mice. Two-way ANOVA with two-stage step-up correction. **b,** Chronic colitis induced by three 7-day cycles of 2% DSS in bone-marrow chimeras (BMC) generated using a 1:1 mixture of CD45.1 wild-type bone marrow (*n* = 5) and CD45.2 CCR7^WT^ or CCR7^ΔcDC^ bone marrow (*n* = 5), with longitudinal body weight and colon length. Two-way ANOVA with two-stage step-up correction for body weight. Unpaired t test with Welch’s correction for colon length. **c,** Gating strategy for flow-cytometric analysis of T cells in mouse colon. **d,** Flow-cytometric analysis of colonic CD4+ Tconv cells from CCR7^WT^ (*n* = 5 vehicle, 5 DSS) and CCR7^ΔcDC^ mice (*n* = 4 vehicle, 5 DSS) in untreated mice and mice at day 7 after DSS treatment. One-way ANOVA with two-stage step-up correction. **e,** FOXP3+ Treg cells as a proportion of CD3+ T cells within lymphoid aggregates and in adjacent or distal tissue from CCR7^WT^ (*n* = 7 aggregates, 13 tissue regions next to aggregates and 11 tissue regions far from the aggregates from 3 mice) and CCR7^ΔcDC^ mice (*n* = 11 aggregates, 17 tissue regions next to the aggregates and 16 tissue regions far from the aggregates from 4 mice) at day 7 after DSS treatment. One-way ANOVA with Fisher’s LSD test. **f,** Flow-cytometric analysis of colonic CD4+ Tconv cells from CCR7^WT^ (*n* = 5 untreated DSS, 4 treated DSS) and CCR7^ΔcDC^ (*n* = 4 untreated DSS, 5 treated DSS) mice at day 7 after DSS at day 7 after DSS treatment with or without FLT3L–Fc. One-way ANOVA with two-stage step-up correction. Data are mean ± s.e.m.

## References

1. Sato, Y., Silina, K., Van Den Broek, M., Hirahara, K. & Yanagita, M. The roles of tertiary lymphoid structures in chronic diseases. Nat. Rev. Nephrol. 19, (2023).

2. Taniguchi, K., Yoshikawa, T. & Yanagita, M. The roles of tertiary lymphoid structures in orchestrating immune responses in peripheral organs. Inflamm. Regen. 45, 28 (2025).

3. Reste, M. et al. The role of dendritic cells in tertiary lymphoid structures: implications in cancer and autoimmune diseases. Front. Immunol. 15, (2024).

4. Schumacher, T. N. & Thommen, D. S. Tertiary lymphoid structures in cancer. Science 375, (2022).

5. Cabrita, R. et al. Tertiary lymphoid structures improve immunotherapy and survival in melanoma. Nature 577, (2020).

6. Helmink, B. A. et al. B cells and tertiary lymphoid structures promote immunotherapy response. Nature 577, (2020).

7. Corsiero, E. et al. Autoimmunity to stromal-derived autoantigens in rheumatoid ectopic germinal centers exacerbates arthritis and affects clinical response. J. Clin. Invest. 134, (2024).

8. Koscsó, B. et al. Gut-resident CX3CR1^hi^ macrophages induce tertiary lymphoid structures and IgA response in situ. Sci. Immunol. 5, (2020).

9. James, K. R. et al. Distinct microbial and immune niches of the human colon. Nat. Immunol. 21, (2020).

10. Sato, Y. et al. Developmental stages of tertiary lymphoid tissue reflect local injury and inflammation in mouse and human kidneys. Kidney Int. 98, (2020).

11. Toriu, N. et al. Aligning cellular and molecular components in age-dependent tertiary lymphoid tissues of kidney and liver. PLOS ONE 20, (2025).

12. Ciccimarra, R. et al. Histological characterization and spatial profiling of age-induced tertiary lymphoid structures in mouse lung parenchyma. Ann. Anat. - Anat. Anz. 260, 152660 (2025).

13. Andersson, R., Mejia-Ramirez, E. & Florian, M. C. Haematopoietic ageing in health and lifespan. Nat. Cell Biol. 27, 1398–1410 (2025).

14. López-Otín, C., Blasco, M. A., Partridge, L., Serrano, M. & Kroemer, G. Hallmarks of aging: An expanding universe. Cell 186, (2023).

15. Kataru, R. P. et al. Structural and Functional Changes in Aged Skin Lymphatic Vessels. *Front*. Aging 3, 864860 (2022).

16. Zolla, V. et al. Aging-related anatomical and biochemical changes in lymphatic collectors impair lymph transport, fluid homeostasis, and pathogen clearance. Aging Cell 14, 582–594 (2015).

17. Lei, P.-J. et al. Aging-induced changes in lymphatic muscle cell transcriptomes are associated with reduced pumping of peripheral collecting lymphatic vessels in mice. Dev. Cell 60, 1118–1133.e5 (2025).

18. Wang, B., Han, J., Elisseeff, J. H. & Demaria, M. The senescence-associated secretory phenotype and its physiological and pathological implications. Nat. Rev. Mol. Cell Biol. 25, 958–978 (2024).

19. Cabeza-Cabrerizo, M., Cardoso, A., Minutti, C. M., Pereira Da Costa, M. & Reis E Sousa, C. Dendritic Cells Revisited. Annu. Rev. Immunol. 39, (2021).

20. Ohl, L. et al. CCR7 Governs Skin Dendritic Cell Migration under Inflammatory and Steady-State Conditions. Immunity 21, 279–288 (2004).

21. Dai, H. et al. Attenuating age-related decline in dendritic cell migration improves vaccine efficacy via gut-immune crosstalk. *Nat*. Aging 6, 816–830 (2026).

22. Grolleau-Julius, A., Harning, E. K., Abernathy, L. M. & Yung, R. L. Impaired Dendritic Cell Function in Aging Leads to Defective Antitumor Immunity. Cancer Res. 68, 6341–6349 (2008).

23. Sikkema, L. et al. An integrated cell atlas of the lung in health and disease. Nat. Med. 29, (2023).

24. Schraml, B. U. et al. Genetic Tracing via DNGR-1 Expression History Defines Dendritic Cells as a Hematopoietic Lineage. Cell 154, 843–858 (2013).

25. Davalos-Misslitz, A. C. M., et al. Generalized multi-organ autoimmunity in CCR7-deficient mice. Eur. J. Immunol. 37, 613–622 (2007).

26. Nitta, T., Nitta, S., Lei, Y., Lipp, M. & Takahama, Y. CCR7-mediated migration of developing thymocytes to the medulla is essential for negative selection to tissue-restricted antigens. Proc. Natl. Acad. Sci. 106, 17129–17133 (2009).

27. Decalf, J. et al. Effectorless Fc-fusion improves FLT3L drug-like properties for cancer immunotherapy combinations. eBioMedicine 118, (2025).

28. Mattiuz, R. et al. Dendritic cells control tertiary lymphoid structure development and maintenance in cancer. Science 393, (2026).

29. Debacq-Chainiaux, F., Erusalimsky, J. D., Campisi, J. & Toussaint, O. Protocols to detect senescence-associated beta-galactosidase (SA-βgal) activity, a biomarker of senescent cells in culture and in vivo. Nat. Protoc. 4, 1798–1806 (2009).

30. Robinson, M. D., McCarthy, D. J. & Smyth, G. K. edgeR : a Bioconductor package for differential expression analysis of digital gene expression data. Bioinformatics 26, 139–140 (2010).

31. Ritchie, M. E. et al. limma powers differential expression analyses for RNA-sequencing and microarray studies. Nucleic Acids Res. 43, (2015).

32. Liu, R. et al. Why weight? Modelling sample and observational level variability improves power in RNA-seq analyses. Nucleic Acids Res. 43, e97–e97 (2015).

33. Phipson, B., Lee, S., Majewski, I. J., Alexander, W. S. & Smyth, G. K. Robust hyperparameter estimation protects against hypervariable genes and improves power to detect differential expression. Ann. Appl. Stat. 10, (2016).

34. Wickham, H. *Ggplot2*. (Springer International Publishing, Cham, 2016). doi:10.1007/978-3-319-24277-4.

35. Veličković, M. et al. Coupling microdroplet-based sample preparation, multiplexed isobaric labeling, and nanoflow peptide fractionation for deep proteome profiling of tissue microenvironment. Preprint at 10.1101/2023.03.13.531822 (2023).

36. Zhu, Y. et al. Spatially Resolved Proteome Mapping of Laser Capture Microdissected Tissue with Automated Sample Transfer to Nanodroplets. Mol. Cell. Proteomics 17, 1864–1874 (2018).

